# *In silico* network-based screening reveals candidates for endothelial dysfunction therapy

**DOI:** 10.1101/2022.11.17.516953

**Authors:** Iguaracy Pinheiro-de-Sousa, Girolamo Giudice, Miriam Helena Fonseca-Alaniz, Silvestre Massimo Modestia, Sarah Viana Mattioli, Yun Fang, Evangelia Petsalaki, Jose E. Krieger

**Author notes:** correspondence should be addressed to Evangelia Petsalaki and Jose E. Krieger.

## Abstract

Endothelial dysfunction (ED) is a hallmark of cardiovascular (CV) disorders and influences their progression; however, there are currently no direct therapeutic targets, primarily due to the lack of knowledge regarding ED’s molecular basis. We used a computational approach to identify candidate targets for ED treatment. We constructed an ED disease gene network by combining the integration of epigenomics (ATAC-seq and ChIP-seq-H3K27ac) and transcriptomics data (RNA-seq) from human aorta endothelial cells (HAEC) exposed to surrogates of primary CV risk factors using network propagation. We then used *in silico* perturbation to prioritise genes that could influence the ED network most when removed. This process resulted in identifying 17 key candidates for which chemical inhibitors are available. These are genes associated with ED and atherosclerosis, and drugs that target those genes have not yet been tested for the treatment of CV disorders. The EGLN3 target and its inhibitor displayed significant anti-inflammatory and antioxidant properties in ECs assessed using a high-content screening platform. These findings illustrate the potential of *in silico* knockouts to discover disease-specific candidate targets for drug development or repositioning.

## INTRODUCTION

The endothelium is a monolayer lining of blood and lymphatic vessels (Boulanger, 2016; Lüscher and Barton, 1997) that plays a critical role in maintaining homeostasis by regulating the vascular tone and angiogenesis while promoting an antioxidant, anti-inflammatory, and anti-thrombogenic response (Fang, Wu, and Birukov, 2019; Godo and Shimokawa, 2017). It is distributed across the entire body and is therefore exposed to various pressure and flow patterns. Regions of arterial branching and curvature, which are also the predominant location for the development of atherosclerotic plaques (Liu et al., 2021; Peiffer, Sherwin, and Weinberg, 2013; Souilhol et al., 2020), are exposed to disturbed shear forces and display early signs of endothelial dysfunction (ED) (Liu et al., 2021; Mudau et al., 2012; Peiffer et al., 2013). These disturbed shear forces are combined with cardiovascular risk factors such as high cholesterol, induce chronic inflammation, and accelerate ED and atherosclerosis (Gencer et al., 2021; Steinberg and Witztum, 2010; de Vries and Quax, 2016).

Historically, the definition of ED included only the reduced capacity of regulating vascular tone (reduced nitric oxide [NO] production). However, it has recently been expanded to include increased oxidative stress and chronic inflammation, compromised barrier integrity, injury repair, and other functions (Xu et al., 2021). ED is associated with most cardiovascular diseases (CVDs) and is a prognostic biomarker for future cardiovascular events such as myocardial infarction or stroke (Cyr et al., 2020; Deanfield, Halcox, and Rabelink, 2007; Mudau et al., 2012). Not surprisingly, due to the multi-functional role of endothelial cells (ECs) and clinical relevance for many CVDs, ED has been considered a potential therapeutic target for several vascular disorders (Daiber et al., 2017; Huynh and Heo 2019; Premer et al., 2019). Nevertheless, despite the importance of ED to human health, the underlying causal mechanisms remain unknown, which hinders the development of therapies directly targeting ED.

ED is typically associated with multiple cardiovascular risk factors and genetic susceptibility rather than single perturbations (Daiber and Chlopicki 2020). Recently, we showed that individual risk factors hierarchically influence endothelial gene expression, and a complex regulation emerges when cardiovascular risk factors are clustered, precluding anticipation of the overall response based on the summation of the individual effects (Pinheiro-de-Sousa et al., 2022). This finding suggests that system biology approaches can integrate various layers of biological information to dissect dysregulated pathways and prioritise core ED-driving genes.

Network-based approaches that combine integrated network assembly with *in silico* perturbations or gene network corrections have been successful in identifying targets for heart valve disease (Theodoris et al., 2020), cirrhosis (Zaffaroni et al., 2019) and other diseases (Cava and Castiglioni, 2019). These approaches typically use executable models (Davidich and Bornholdt 2013), network propagation (Santolini and Barabási 2018) or efficiency analysis (Cava and Castiglioni 2019) on a specific network. Their limitation is that they don’t consider changes in the wiring of this network as a result of this perturbation but rather identify potentially affected nodes from a perturbation. Theodoris et al, recently acquired data upon perturbation and then formed perturbed networks which were then used in a machine learning model to evaluate whether the perturbations return a disease network to the ‘healthy state’ (Theodoris et al. 2020). However, this approach is costly and time consuming as it requires extensive additional data collection. A recent method combined identification of the most relevant transcription factors in a gene regulatory network with identification of upstream signalling molecules that can regulate them (Zaffaroni, Okawa, Morales-Ruiz, and del Sol 2019). While this circumvents several of the limitations described above, it is limited to 75 annotated signalling pathways that include only ∼2,500 nodes.

Here, we combined multi-omics data integration with network inference and a new strategy for *in silico* perturbations to identify and validate targets and drugs that can be repurposed for ED (Figure 1).

**Figure 1.**
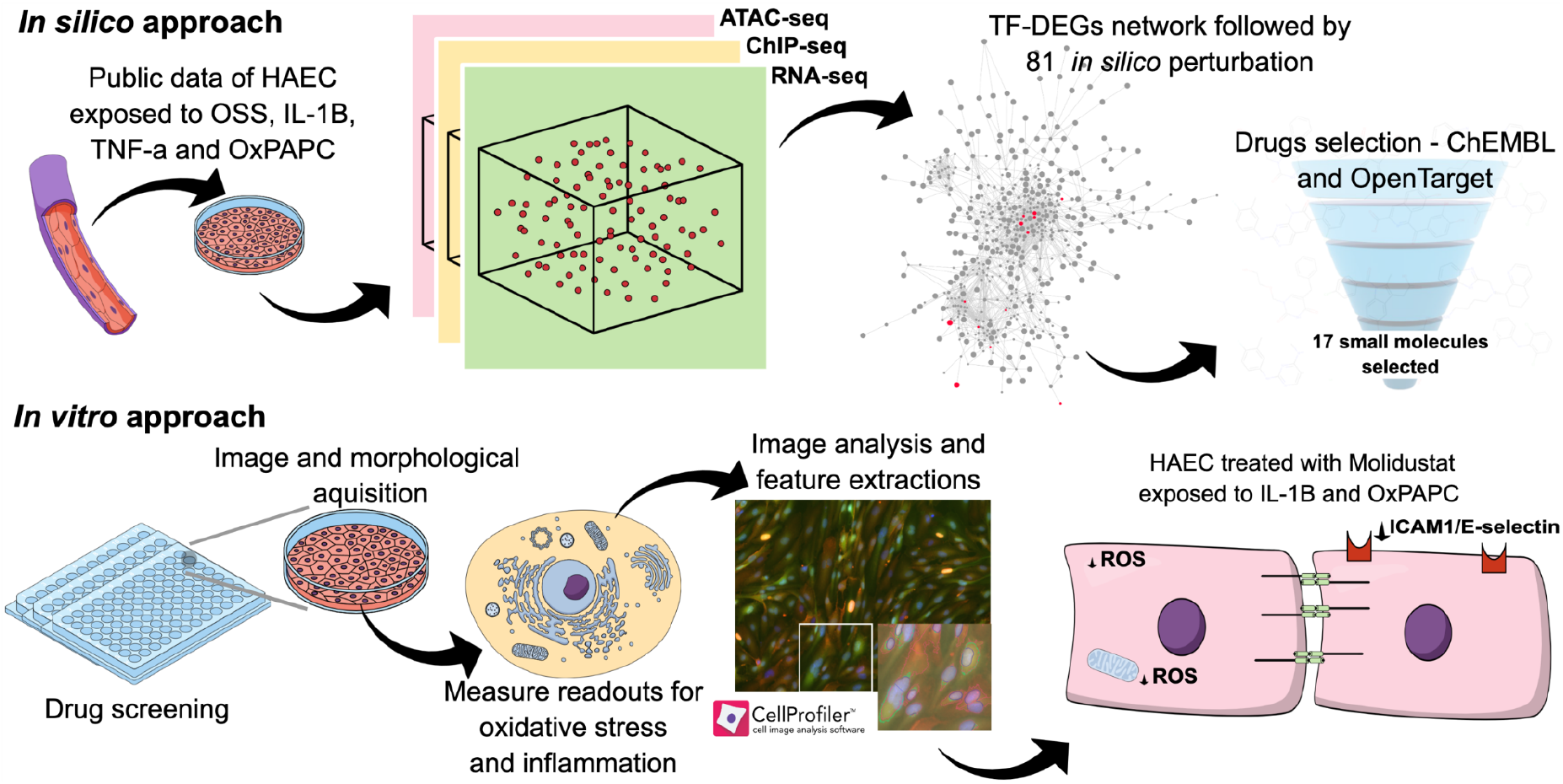
Schematic overview of the integrative multi-omics network analysis to identify ED targets. Multi-omics (ATAC-Seq, H3K27ac ChIP-Seq, and RNA-Seq) data from human aorta endothelial cells (HAEC) exposed to surrogates of CV risk factors were integrated. These surrogates included mechanical force oscillatory shear stress (OSS) and chemical stress with TNF-α, IL-1β, and OxPAPC (Methods). We built an ED disease regulatory network using the transcription factors (TFs) enriched in the common regulatory regions of all stimuli and their downstream differentially expressed genes (DEGs). TF-DEGs targeted with drugs available were knocked out in silico to identify genes with the most disruptive effect in the disease network. A subset of those targets with available drugs was further screened in vitro in HAECs using immunofluorescence assays and image analysis to measure reactive oxygen species (ROS) and inflammatory markers (ICAM1).

## RESULTS

### Surrogates of cardiovascular risk factors OSS, IL-1β, TNF-α, and OxPAPC have common effects on the epigenetic landscape and transcriptional programme

We hypothesised that combining data collected using various surrogates of CV disease risk factors would allow the extraction of more relevant gene regulatory regions specific to ED. Therefore, we re-analysed public ChIP-seq, ATAC-seq, and RNA-seq data measured from HAECs exposed to either mechanical stimuli such as laminar (LSS) and oscillatory shear stress (OSS) (Krause et al., 2018) or chemical stimuli including the inflammatory cytokines IL-1β and TNF-α and oxidised phospholipids (OxPAPC) (Hogan et al., 2017). Chromatin accessibility, measured by ATAC-seq analysis, identified 17,483 open chromatin regions common among HAECs subjected to OSS, IL-1β, TNF-α, and OxPAPC (Figure 2A and S1A). ChIP-seq data analysis revealed that 14,149 enhancer-like elements, marked by significant levels of acetylation of lysine 27 (H3K27ac), were common among the stimuli (Figure 2A and S1B). ATAC-seq peaks were used to centre the enhancer-like regions (ChiP-seq peaks), which resulted in 6,630 high-confidence regulatory regions commonly modulated by all stimuli (Figure 2A and S1C). Most of these peaks (∼60%) were found in promoter regions (Figure 2B).

**Figure 2.**
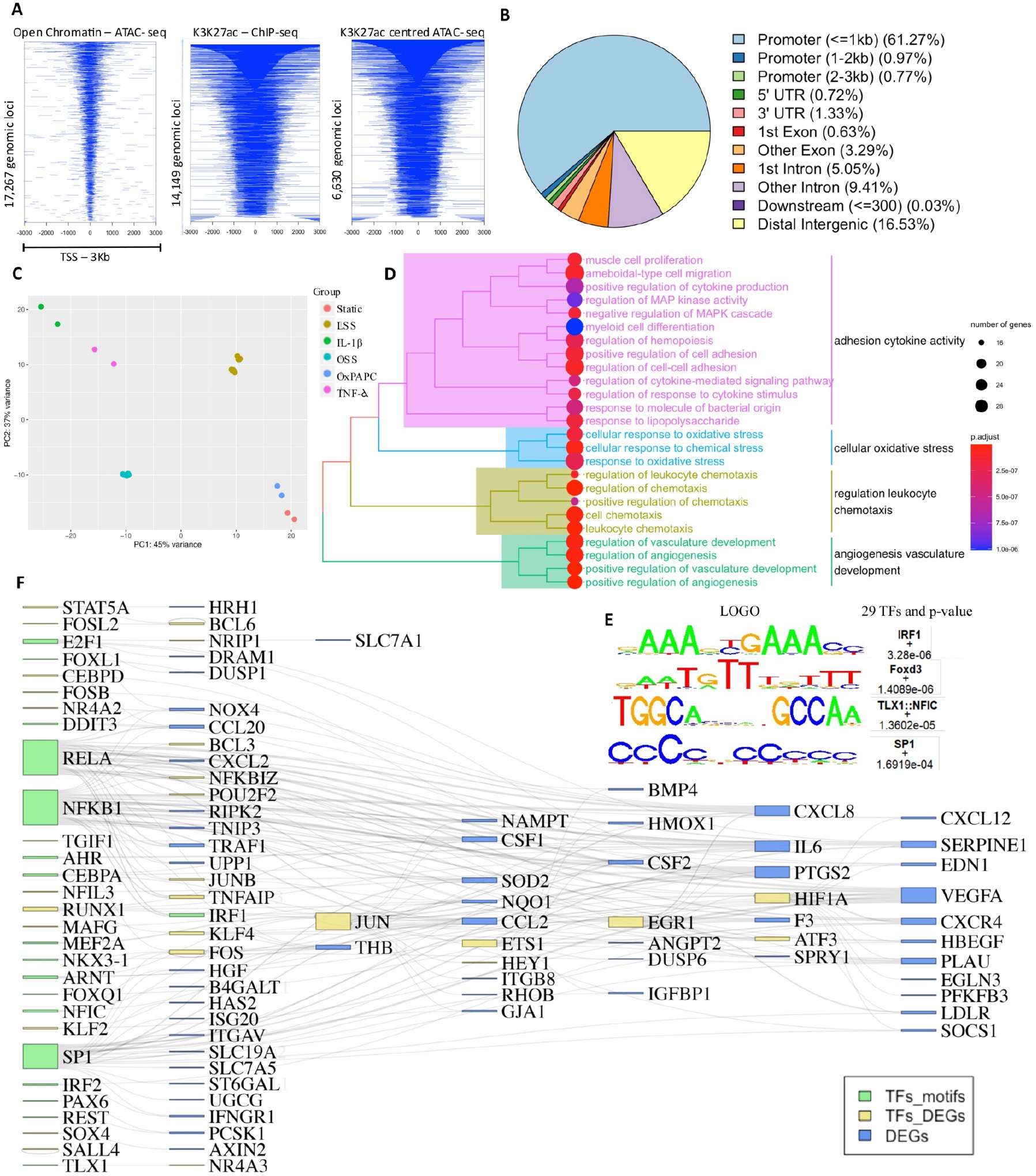
Surrogates of CV risk factors OSS, IL-1β, TNF-α, and OxPAPC modulate common regulatory regions on the genome. **(A)** A heatmap of 17,267 common accessible chromatin regions (ATAC-seq), 14,149 common enhancer-like regions H3K27ac (ChIP-seq) and 6,630 common enhancer-like regions centred on accessible chromatin (<= 3 kb from promoters). **(B)** Distribution of the genomic features of the 6,630 common enhancer-like regions. The genomic features include whether a peak is in the TSS, Exon, 5’ UTR, 3’ UTR, intronic, or intergenic. **(C)** Principal component analysis of gene expression profiles from each sample under IL-1β, TNF-a, and OSS demonstrates the major components PC1 of the variance to separate samples from basal control, OxPAPC, and LSS. PC1 and PC2 accounted for 45% and 37% of the total variance, respectively. **(D)** Gene Ontology (GO) enrichment analysis for biological processes (BP) followed by redundancy term reduction analysis (adjusted p ≤ 0.05). We considered 356 DEGs (adjusted p ≤ 0.05 and log2 |fold change| ≥ 1.5) for the GO enrichment analysis identified under the 6,630 peaks. **(E)** The top four enriched motifs in the centre of enhancer-like regions, along with the relevant transcription factor family, motif sequence and enrichment log p-values. Enrichment was calculated from a 200 bp sequence centred on chromatin accessibility. **(F)** Sankey diagram showing the TF-DEGs interactions.

The transcriptomics data revealed the differences and hierarchies among the stimuli. Our principal component analysis showed that 45% of the source of variation in gene expression is explained by the PC1, which separates LSS, static basal control, and OxPAPC from the other stimuli, while PC2 clustered OSS, static basal control, and OxPAPC from LSS, IL-1β, and TNF-α (Figure 2C). Using hierarchical clustering, we identified two major groups, one including IL-1β, TNF-α, and OSS and the other including LSS, static basal control, and OxPAPC (Figure S1D). Our differential expression analysis considered the comparisons of OSS vs LSS, whereas IL-1β, TNF-α and OxPAPC treatments were compared to their static control owing to data availability. We found 578 (OSS) and 928 (IL-1β), 594 (TNF-α), and 186 (OxPAPC) DEGs in the respective cardiovascular risk factor models. Because we were interested in the common genes and pathways to these risk factors, we used the genes annotated to the 6,630 peaks that were differentially expressed in at least one condition, resulting in 356 DEGs. Known dysregulated ED processes were enriched in this set of genes, including inflammatory pathways (e.g., cytokine production, cell-adhesion molecules, and leukocyte chemotaxis), oxidative stress, and regulation of angiogenesis and vasculature development (Figure 2D).

To identify the gene regulatory network and processes relevant to ED, we then identified key transcription factors (TFs) relevant to the abovementioned stimuli and their downstream DEGs. We identified seven significantly enriched motifs by performing *de novo* motif enrichment analysis at the 6,630 peaks. We considered only the significant TFs predicted to bind these motifs (p < 0.05, Figure S1E; Methods)—29 in total. The most significant were Foxd3 (p = 3.3e-06), IRF1 (p = 1.4e-06), TLX1: NFIC (p = 1.4e-05), and SP1 (p = 1.6919e-06) (Figure 2E). The TF-DEG interactions were considered only when the DEGs were present at the 6,630 enhancer-like regions, and these interactions were supported by the TRRUST database (Han et al., 2018). The resulting TF-DEGs interactions comprise 100 genes (Figure 2F, Table S1), of which 46 are TFs (18 motifs enriched, 28 differentially expressed, Figure 2F) and 54 are not (in blue, Figure 2F). Among the TF-DEGs identified above, NFKB1 and RELA, associated with inflammatory response (Mussbacher et al., 2019), regulate the largest amount of downstream DEGs. Moreover, NFKB1 is upregulated in the IL-1β and TNF-α conditions compared to their basal control.

### Network propagation reveals an ED network that is enriched in DEGs from human ECs located at human atherosclerotic plaques

We then used a network propagation strategy to identify the processes and network regions most affected by the deregulation of these ED genes. To this end, we applied the random-walk-with-restart (RWR) algorithm (Tong, Faloutsos, and Pan, 2008) to a human protein interaction network (Hornbeck et al., 2015; Kerrien et al., 2012; Surdo et al., 2017; Türei, Korcsmáros, and Saez-Rodriguez 2016), using the 86 genes among them as seeds that were differentially expressed in at least one condition (Figure 3A, Methods). The resulting ED network comprised 838 nodes and 1,991 edges (Figure S2A, Table S2). Functional enrichment analysis showed enriched inflammatory pathways among TNF, IL-1 response, and cytokine-mediated cascade (Figure S2B), boosting the inflammatory response signal we observed for the biological process enriched for the 356 DEGs under the 6,630 peaks (Figure 2D).

**Figure 3.**
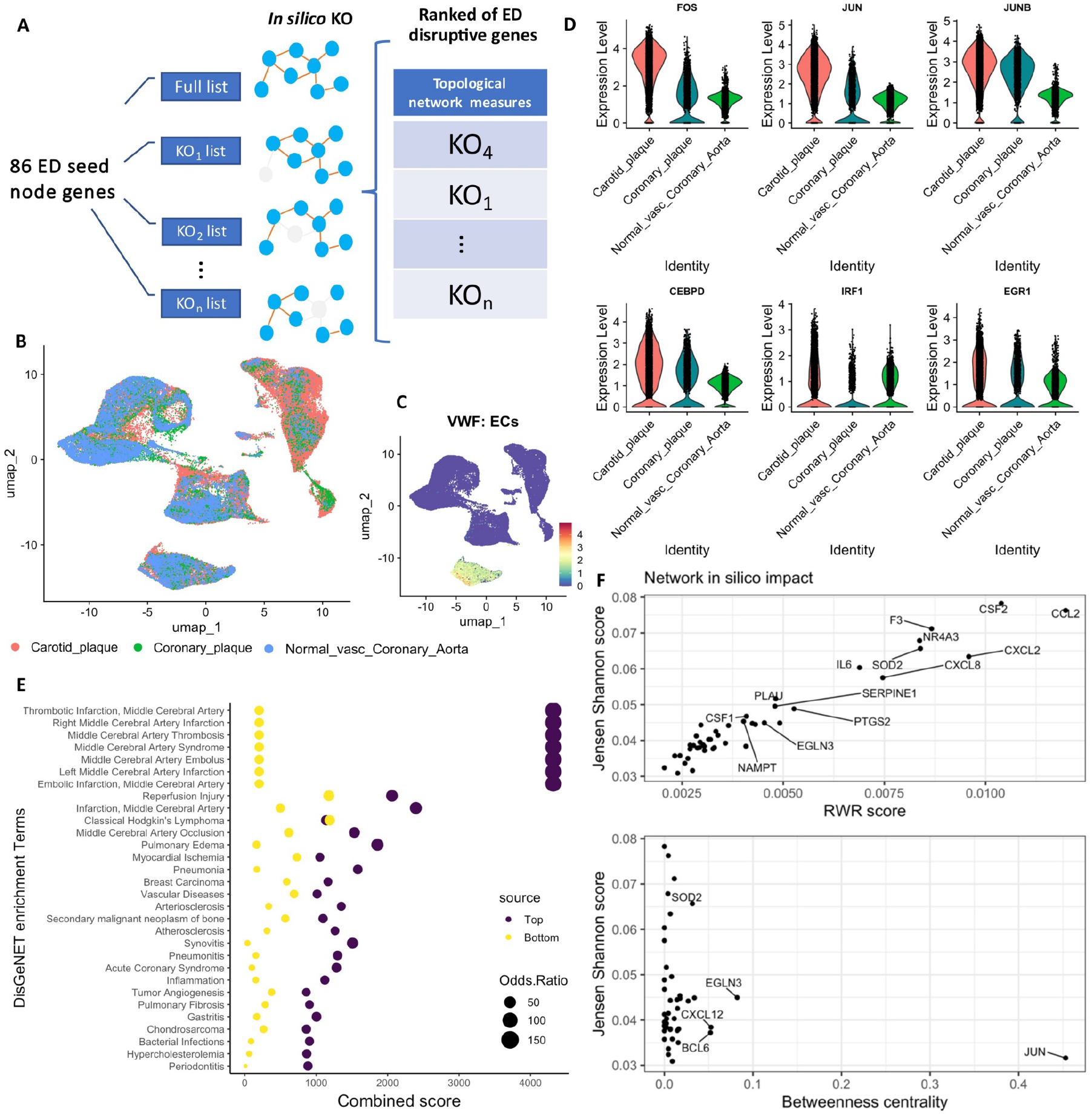
in silico knockout (KO) screen identifies 17 genes that present a disruptive effect in the ED disease network. **(A)** Schematic process of generating the ED disease network and the ranking of in silico KO perturbations with the most disruptive effect in the network (Methods). **(B)** Uniform manifold approximation and projection (UMAP) visualisation of the integrated scRNAseq datasets used to validate the ED disease network (n = 69,385 cells). **(C)** UMAP visualisation of a critical endothelial cell marker VWF **(D)** Violin plot of the TFs differentially expressed in vitro (HAECs under the stimuli) and in vivo (upregulated in ECs at the atherosclerotic plaque). **(F)** Disease enrichment analysis of the 81 KO genes ranked by the Jensen-Shannon distance showing the enriched terms of the top (41 genes) vs bottom (40 genes) **(E)** Network topology measurements used to rank the disruptive effect of the in-silico KOs.

To confirm whether our ED network represented relevant processes *in vivo*, we re-analysed three scRNA-seq datasets from human atherosclerotic vasculature (Alsaigh et al., 2020; Jones et al., 2022; Wirka et al., 2019) (Figure S2C). Two atherosclerotic plaque datasets were compared to those acquired from vasculature without plaques (Figure 3B). ECs were further subdivided (Figure 3C), and differential expression analysis in ECs from the plaque vs normal vasculature identified 705 DEGs (526 upregulated and 179 downregulated, Table S3). Among the 526 upregulated DEGs, the top genes are known to be associated with atherosclerosis plaque, including matrix Gla protein (Herrmann et al., 2000), VIM (Shi, Sun, and Kaptoge, 2021), and B2M (Shi et al., 2021) (Figure S2D). The DEGs identified in the ECs plaques were significantly enriched in our ED network (hypergeometric test p < 0.008) compared to the background human protein interaction network (Figure S2E), suggesting that our ED network is relevant for *in vivo* systems. Interestingly, 11 of the 28 differentially expressed TFs (in yellow, Figure 2F) were also differentially expressed in *in vivo* ECs (ten in the same direction, upregulated in the atherosclerotic plaque), suggesting that they might be among the primary regulators of the downstream effect in response to CV risk factors (Figure 3D, Table S1).

### *In silico* knockout (KO) perturbations identified a set of putative gene targets for ED

We hypothesised that genes that disrupt the ED network topology when perturbed are critical for targeting ED. To this end, we performed *in silico* KO perturbations (Figure 3A).

We hypothesised that genes critical for maintaining the topology of the ED network could be good targets for the condition. Therefore, we removed each of the 81 differentially expressed ED genes that were upregulated among the initial 86, repeated the RWR process, and evaluated the change in the network topology using the Jensen-Shannon distance (Endres and Schindelin 2003) between networks and other topological parameters (Table S4, Methods). Comparing the functional disease enrichment of the top and bottom-ranked genes supported our hypothesis and ranking because the top-ranked genes were significantly more enriched in CVD pathways such as thrombotic infarction related to the cerebral artery, myocardial ischemia, atherosclerosis, and others (Figure 3E).

We then identified potential drugs for repurposing targeting our top-ranked genes; 44 of the 81 initial seed nodes were annotated as targeted by drugs in ChEMBL and the OpenTarget databases (Ochoa et al., 2021; Willighagen et al., 2013). Several of these were critical for the ED network, including *CCL2, CSF2, CXCL12, F3, CXCL8, IL6, SOD2*, and *NR4A3*, among the highest-scoring ones (Figure 3F above). Considering the betweenness centrality measure (Yoon, Blumer, and Lee, 2006), we found *JUN, EGLN3, CXCL12*, and *BCL6* to be among the most important for joining dense subnetworks within our ED disease network (Figure 3F bottom). The DisGeNet (Piñero et al., 2020) score (Table S4, Methods) showed that several of these genes had been previously associated with ED or CVDs and inflammatory pathways (*CSF2, CXCL8/12, CXCR4, CCL2*, and *IL6*) (Galkina and Ley, 2009; Gencer et al., 2021). However, for some (e.g., *EGLN3*), their role in ED remains poorly understood (Fu and Taubman, 2013; Gerber et al., 2009) (Figure 3F).

### The EGLN3 inhibitor is a candidate for drug repurposing against ED

To confirm whether the genes critical for the ED network are also potential targets for ED, we created an *in vitro* validation strategy.

First, we filtered the drugs available (OpenTarget and ChEMBL) that target those genes based on drug-like properties (Methods), resulting in the selection of one drug for each of the 17 top-scoring genes (Figure 3A; Table S5).

ROS are generated when ECs are exposed to factors such as OxLDL/OxPAPC that can lead to excessive endothelial permeability and reduction of NO availability (Feng et al., 2014; Liu et al., 2014; Springstead et al., 2012). Inflammation also participates in the initiation of ED and the progression of atherosclerosis. Among the several chemokines and interferons produced in this process, the inflammatory marker intercellular adhesion molecule-1 (ICAM1) is a critical molecule expressed in inflamed ECs that allows monocyte adhesion and plaque growth (Fisker Hag, Pedersen, and Kjaer, 2008; Gencer et al., 2021; Ley et al., 2007; Sucosky et al., 2009). Therefore, we performed image-based high-content screening (Lin et al., 2020; Mattiazzi Usaj et al., 2016) using the 17 drugs in HAEC in the presence of the combined stimuli of IL-1β (10 ng/mL) and OxPAPC (50 µg/mL) and quantified the changes of these essential readouts for ED (i.e., ROS and protein expression of ICAM1 [Figures 1 and 4]) as a proxy for EC function/dysfunction.

**Figure 4.**
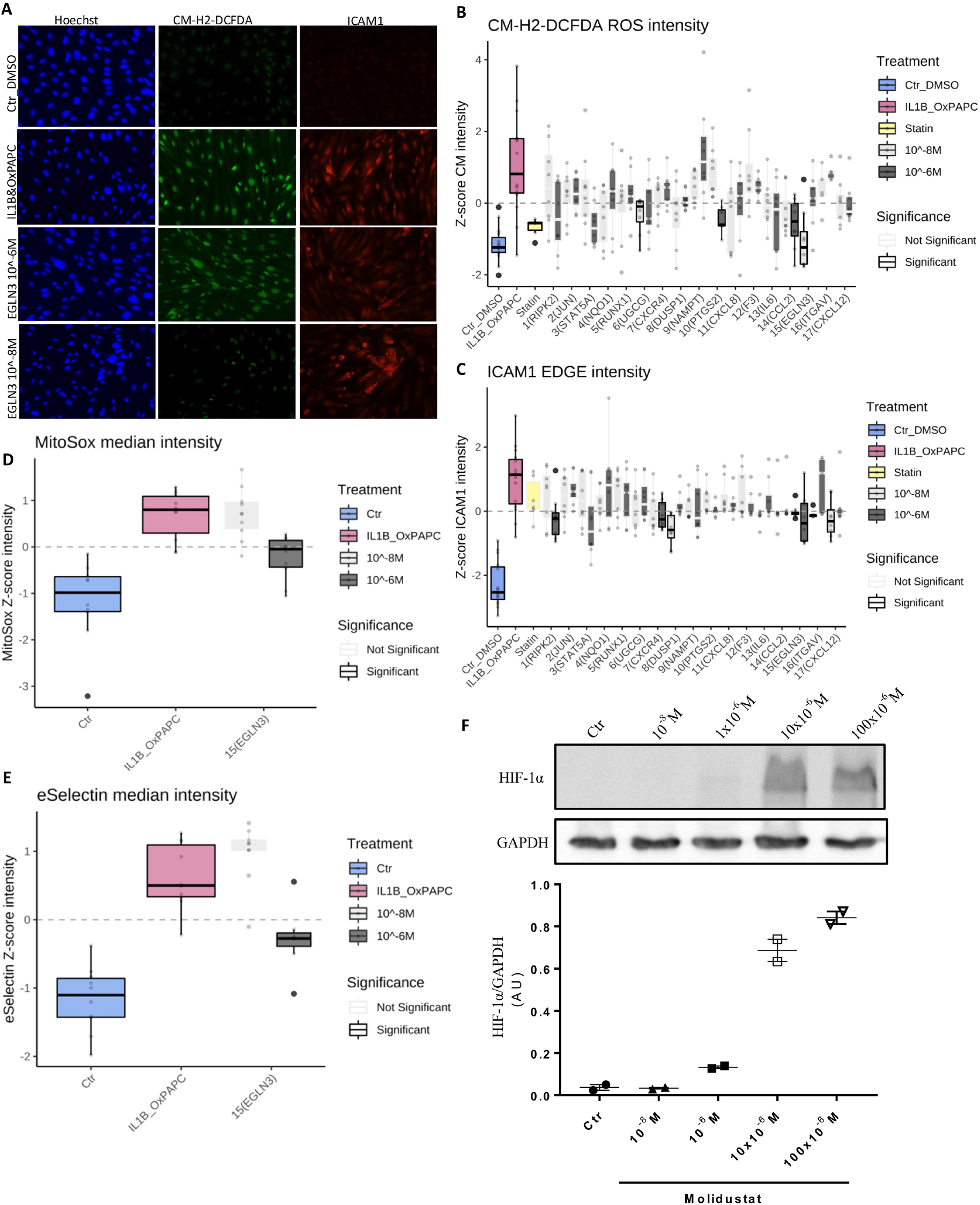
Inhibition of the UGCG, PTGS2, CCL2, and EGLN3 proteins showed significant antioxidant effects while inhibiting RIPK2, CXCR4, DUSP1, EGLN3, and CXCL12 displayed an anti-inflammatory effect in HAECs under IL-1 and OxPAPC stimuli. **(A)** Representative fluorescent images showing the nuclei (blue), ROS (green), and ICAM1 (red) staining of HAEC under DMSO, IL-1β, and OxPAPC, and treated with EGLN3 inhibitor (Molidustat, 10^−8^ M and 10^−6^ M). **(B, C)** The plot shows signal intensity from ROS (probe CM-H2-DCFDA, green) and ICAM1 (antibody anti-ICAM1, red). **(D)** Mitochondrial ROS (MitoSox probe) intensity only for the EGLN3 inhibitor. **(E)** E-selectin inflammatory marker (antibody anti-E-selectin) intensity only for the EGLN3 inhibitor. Signal intensity was normalised to the internal control DMSO in each plate, followed by z-score normalisation across plates. P-values were computed by multiple pairwise comparisons with the Wilcon test followed by Benjamini–Hochberg correction (BH), comparing all conditions vs HAECs treated with IL1B&OxPAPC (positive disease control). We considered a significant p-adjusted value < 0.05 (coloured in the plot). Data are presented as mean ± standard deviation. All experiments were repeated independently at least twice, with technical replicates from four to five wells by condition. **(F)** HIF-1α was determined by western blot in HAEC treated with Molidustat (EGLN3 inhibitor) with doses from 10^−8^ M to 100×10^−6^ M for 48 h. GAPDH served as a housekeeping protein. Two biological replicates are presented.

First, we performed a cell viability test for two concentrations of the drugs (10^−6^ M and 10^−8^ M). Neither drug nor the dimethyl sulfoxide (DMSO) control at either concentration was cytotoxic, compared to cells at 24 h of starving (Figure S3A and B). ROS levels and ICAM1 expression were also measured for all drugs in HAECs under basal conditions for 48 h (without IL-1β and OxPAPC stimuli). The inhibition of IL6, CCL2, and EGLN3 suppressed basal ROS levels production, while the inhibition of RUNX1, CXCR4, and NAMPT increased ROS levels compared to the DMSO control (Figure S3C). ICAM1 signal intensity was significantly decreased for STAT5A, UGCG, CXCR4, IL6, and EGLN3 when inhibited in HAECs under basal conditions (Figure S3D).

We next treated HAECs with IL-1β and OxPAPC and found more significant expression of ICAM1 and production of ROS (Figure 4A) than the DMSO control. We used atorvastatin (5 µM) as a positive drug control because it is widely used for patients at high CVD risk and displays a pleiotropic beneficial effect on endothelial function (Oesterle, Laufs, and Liao, 2017; Stancu and Sima, 2001). The inhibition of UGCG, PTGS2, CCL2, and EGLN3 suppressed ROS production (Figure 4B). The inhibition of RIPK2, CXCR4, DUSP1, EGLN3, and CXCL12 suppressed ICAM1 protein expression in HAEC treated with IL-1β and OxPAPC (Figure 4C). Atorvastatin presented an antioxidant effect, suppressing ROS levels as expected (Oesterle et al., 2017) but did not show changes in ICAM1 expression compared to the combined stimulus of IL-1β and OxPAPC. The drug that inhibits the EGLN3 protein was unique in displaying antioxidant and anti-inflammatory responses in HAECs exposed to IL-1β and OxPAPC. To further investigate the role of EGLN3 inhibition in ROS and inflammatory responses, we measured mitochondrial ROS and E-selectin, another inflammatory molecule increased in inflamed ECs (Chiu et al., 2004; Gencer et al., 2021). The inhibition of EGLN3 suppressed the mitochondrial ROS and E-selectin production in HAEC under IL-1β and OxPAPC treatment (Figures 4D and E), corroborating the antioxidant and anti-inflammatory potential of this drug in ECs.

The EGLN3 protein (prolyl-hydroxylase Egl nine homolog 3), also known as PHD3, hydroxylates proline residues in target proteins such as PKM, TELO2, ATF4, and HIF-1α (Appelhoffl et al., 2004; Luo et al., 2011; Xie et al., 2012). PHD inhibitors stabilise HIF-1α by inhibiting its degradation by prolyl-hydroxylase enzymes (Joharapurkar et al., 2018). The PHD inhibitor used here was Molidustat (CAS#1154028-82-6) (Beck et al., 2018). As HIF-1α is the downstream target of this drug, we evaluated its protein expression in the basal condition (without IL-1β and OxPAPC) and demonstrated dose-dependent (10^−8^ M to 100 × 10^−6^ M) enhancement of HIF-1α protein expression compared with the DMSO control (Figure 4F). Next, we measured the gene expression of known targets of HIF-1α and observed increased expression of *ALDOA, LDHA*, and *CXCR4* and decreased expression of *NOS3* in response to Molidustat compared to the DMSO control (Figure S4). These findings confirm the proper functioning of Molidustat.

## DISCUSSION

ED is a hallmark of CVD, and studies have shown that individuals with a dysfunctional endothelium have a higher risk of developing cardiovascular events such as myocardial infarction or stroke (Gokce et al., 2002; Matsuzawa et al., 2013; Schächinger, Britten, and Zeiher, 2000). Nevertheless, most drugs targeting CVDs, such as atherosclerosis or hypertension, have not been developed to specifically target ED (Daiber et al., 2017; Hennigs et al., 2021; Libby, 2021).

Individual cardiovascular risk factors hierarchically influence endothelial gene expression. Combining diverse CVD risk factors results in complex gene network regulation that is not trivial to predict based on the summation of the individual effects (Pinheiro-de-Sousa et al., 2022). Here, we presented a strategy that combines data integration with *in silico* network-based screening to select candidate drugs/gene targets for ED. Considering this complex scenario underlying the pathophysiology of ED, we hypothesised that identifying commonly deregulated genes for multiple surrogates of these risk factors would increase the chance of successfully targeting ED. We first identified 86 genes by integrating ATAC- and Chip-Seq data from HAECs subjected to various surrogates of CV risk factors and intersecting the commonly affected chromatin regions with DEGs identified by RNA-Seq. Using network propagation, these selected genes were then used to generate an ED network. A systematic *in silico* perturbation of the DEGs in the network uncovered the critical genes for the network topology. Among these, we selected 17 targets based on drug availability and properties for experimental evaluation of their effect on ED features (e.g., inflammation and oxidative stress) using relevant primary human ECs and a high-content imaging approach.

Our analysis revealed that the pharmacological inhibition of UGCG, PTGS2, and CCL2 has antioxidant properties, while inhibition of RIPK2, CXCR4, DUSP1, and CXCL12 results in reduced inflammation properties. Molidustat, an inhibitor of the EGLN protein family, had both antioxidant and anti-inflammatory effects.

EGLN3/PHD3 works by hydroxylating proline residues to stabilise proteins such as HIF-1α and inhibit their degradation (Appelhoffl et al., 2004). Activation of HIF-1α has been associated with ED and the formation of atherosclerotic plaques (Abe, Semba, and Takeda, 2017). In ECs at atherosclerotic disturbed flow sites, HIF-1α increases, and its stabilisation promotes ROS via NOX4 (Wu et al., 2017). Treatment with a PHD inhibitor resulted in decreased atherosclerotic plaque accumulation, lower serum cholesterol levels, and reduced blood levels of macrophage aggregates in mice (Rahtu-Korpela et al., 2016). A reduction in myocardial ischaemia-reperfusion injuries was shown after PHD depletion in rats (Neckář et al., 2018; Vogler et al., 2015). According to Necká et al., 2018, ischaemia-reperfusion injuries are associated with high levels of ROS and reduced stability of HIF-1. The inhibition of PHD ameliorates the HIF-1 stabilisation issue during cardiac reperfusion (Neckář et al., 2018).

Here, we demonstrated the accumulation of HIF-1α protein when increasing the dose of Molidustat (Figure 4F). However, the protective effect of this drug was present at lower doses (10^−8^M and 10^−6^M). The HAECs treated with Molidustat cultured under static conditions combined with IL-1β and OxPAPC presented reduced production of generalised and mitochondrial ROS, ICAM1, and E-selectin. Currently, the usage of PHD inhibitors like molidustat to target CVD is limited to pre-clinical animal studies; however, the therapeutic use of PHD inhibitors is in late-stage clinical trials (Joharapurkar et al., 2018; Tanaka and Eckardt, 2018) for the treatment of anaemia associated with chronic diseases (Amit A. Joharapurkar et al., 2018). Therefore, there is a potential chance for repositioning this drug for CVDs in human clinical trials. While the detailed mechanism of action of such drugs warrants further investigation, our perturbed ED networks can provide a starting point for such studies.

Our findings suggest that, by combining the integration of omics datasets from diverse conditions with network analysis, we can identify networks relevant to the common phenotype among these conditions, such as ED, and can help prioritise targets to modulate these phenotypes. Our strategy for *in silico* perturbation simulates the changes in the network topology upon perturbation, by redefining the disease networks upon each virtual KO. By using a global signalling-focused interaction network as the basis of our analysis we can mitigate the bias of signalling focused approaches, while maintaining the advantage of identifying relevant nodes that can modulate the changes in a cell’s state. Nonetheless, while *in silico* KOs are powerful for identifying essential nodes in the network, in general, they are unlikely to predict the direct effect on the cell’s phenotype unless paired with relevant machine-learning approaches. They are thus best suited as a valuable target prioritisation approach for *in vitro* studies and beyond. Nevertheless, using this data-driven approach, we identified eight targets that can ameliorate ED phenotypes *in vitro*, with one of them affecting both phenotypes. Thus, our analysis resulted in the discovery of targets that can be candidates for drug repurposing to ameliorate ED phenotypes and delay or halt CVD progression.

## Data and code availability

All codes developed and used in this study are freely available through: https://zenodo.org/record/7312056#.Y3Z1OOzP0UF

All data used in this study are publicly available with the respective identifiers described in the methods section.

## Author contributions

Conceptualisation: IPS., EP, and JEK;

Data curation: IPS, YF;

Formal analysis: IPS, GG;

Funding acquisition: EP, JEK;

Resources: JEK, EP;

Validation: IPS, MHFA, SVM, SMM;

Writing, reviewing, and editing: IPS, GG, EP, and JEK

## Acknowledgements

IPS received a fellowship from the Coordination for the Improvement of Higher Education Personnel (CAPES 88882.328126/2019-01) to visit EMBL-EBI. JEK is supported by grants from the Sao Paulo Research Foundation Fundação de Amparo a Pesquisa do Estado de São Paulo [FAPESP-INCT—20214/50889-7 and 2013/17368-0] and the Conselho Nacional de Desenvolvimento Científico e Tecnológico—CNPq [INCT—465586/2014-7 and 309179/2013-0]. EP, GG, and IPS were supported by funding provided by EMBL-EBI.

## Materials and methods

### Datasets

To identify the ED genes, we integrated publicly available RNA-Seq, H3K27ac ChIP-Seq, and ATAC-Seq datasets (GSE89970, GSE112340) from cells HAECs. In the GSE112340 dataset (Krause et al., 2018), data from HAECs was acquired either in static conditions or after 24-h unidirectional or disturbed flows. In the GSE89970 (Hogan et al., 2017), HAECs were subjected to inflammatory cytokines TNF-α (10 ng/mL), IL-1B (10 ng/mL) or OxPAPC (40 µg/mL) for 4h.

### ATAC-seq and ChIP-seq analysis

For the ChIP-seq and ATAC-seq data, we used the pre-processed hg19-aligned data using Bowtie2 (Langmead and Salzberg 2012). Peaks were identified using the MACS2 tool (Kim and Dekker 2018). The BED files from multiple replicates were compiled through consensus voting (https://ro-che.info/articles/2018-07-11-chip-seq-consensus), where the peaks considered were the overlapping peaks between replicates (packages in R, *rtracklayer* and *GenomicRanges*) (Lawrence et al., 2013; Lawrence, Gentleman, and Carey 2009). ATAC-Seq and ChIP-Seq consensus peaks were identified and annotated using the *ChIPSeeke*r R package (Yu, Wang, and He 2015).

We integrated the ChIP-seq and ATAC-seq data by identifying all the peaks that overlap in all conditions using the *ChIPpeakAnno* R package (Zhu et al., 2010). We used the findOverlappingPeaks() function, which uses as input the consensus peaks identified previously for each condition. We considered two peaks overlapping when one resided entirely within the second one. We identified transcriptionally active regions by applying the findOverlappingPeaks() function to centre the ChiP peaks onto the open chromatin ATAC peaks regions. Thus, this final merged BED peak file contained regions common to all stimuli and regions of open chromatin and active enhancers.

### Motif enrichment analysis

For *de novo* motif enrichment, we used *rGADEM* and *MotIV* (R packages) to predict active TFs from the merged BED peak file containing a 200 bp sequence. The *rGADEM* package combines a genetic algorithm (GA) with an expectation-maximization (EM) algorithm to identify the statistically significant motifs. We considered the motifs whose scores reached the significance threshold of E < 0.05. Then, we used MotIV, the JASPAR database (Castro-Mondragon et al., 2022), and the motifs identified by *rGADEM* to identify the TFs. The motifMatch() provided the TFs alignment to the input motifs and their p-value, which we kept all TFs statistically significant (p < 0.05).

### RNA-seq analysis and TF-DEGs associations

The RNA-seq samples were subjected to fastQC quality control, where the sequences were trimmed by 13 bp to create the SAM files, later converted to BAM and aligned to hg19 using HTSeq2 (Putri et al., 2022). Principal component analysis, batch correction effect, and differential expression analysis were performed using the DESeq2 R package (Love, Huber, and Anders, 2014). Differential expression analysis was performed for each condition (IL-1β, TNF-α, and OxPAPC) compared to static baseline control, while OSS was compared to LSS. We consider as DEGs those with an adjusted p < 0.05 and |Log2Foldchange| > 1.5. Using the merged BED peak from the ChIP and ATAC overlapping peaks and their annotated genes, we filtered by identifying which DEGs were present in those peaks. We used the TRRUST database (version 2) (Han et al., 2018) to map the TFs identified to their previously published transcription targets. We considered only the TFs and DEGs whose interactions we supported by published evidence (Figure 2D).

### Pathway enrichment analysis and gene ontology (GO)

We performed an enrichment analysis of BP for the 356 DEGs, differentially expressed in at least one condition under the common 6,630 peaks and the 838 nodes representing the ED disease network using the clusterProfiler package (Yu et al., 2012). We used over-representation analysis (Boyle et al., 2004) to identify the primary BP associated with these TF-DEGs. We considered enriched BP with adjusted p < 0.05. Due to high gene ontology BP redundancy, we applied the treeplot() function using ward.D methods hierarchical clustering the enriched terms with high semantic similarity.

## Network analysis

### Empirical network and random network generation

The human protein-protein interaction network (PIN) was downloaded from IntAct (release 234, last update May 2021) (Kerrien et al., 2012; Orchard et al., 2014). Kinase-kinase and kinase-substrate relationships from PhosphoSitePlus (PMID: 25514926) (last updated May 2021) (Hornbeck et al., 2015), OmniPath (PMID: 27898060) (last release May 2021) (Türei et al., 2016), and SIGNOR 2.0 (PMID: 31665520) (last release May 2021) (Surdo et al., 2017) were included into the PIN. Only proteins annotated in Swiss-Prot (PMID: 33237286) and with at least one GO term (last release May 2021) (PMID: 33290552) were retained. The final PIN comprised 16,407 nodes and 238,035 edges. PIN edge weights were modelled according to the Topological Clustering Semantic Similarity (PMID: 21078182) and calculated using the Semantic Measure Library (PMID: 24108186). To associate a p-value to each network node, we generated 1,000 random networks with the configuration model available in the igraph library (method = vl). Because the configuration model creates new random interactions, the edge weights were updated accordingly.

Each network was corrected for the hub bias according to the following equation:

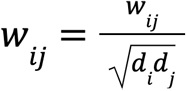

where *W*_*ij*_ indicates the edge weight (i.e., semantic similarity) and *d*_*i*_ and *d*_*j*_ represent the weighted degree of node *I* and node *j*.

### Empirical network propagation

We employed the RWR algorithm (Tong et al., 2008) through the personalized PageRank function available at http://igraph.org to mimic the effect of the gene’s upregulation inside the PIN. For this aim, we used the 81 upregulated DEGs as seed nodes and as weight, their respective log2 fold change normalized and a damping factor equal to 0.7. We called the RWR scores obtained empirical scores.

To assess the significance of the nodes, we repeated the procedure above using the same partition but against the 1,000 random networks generated (see ‘empirical and random network generation’). We then estimated the p-value for each node of the PIN and each partition separately using the following formula:

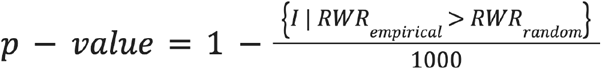

Where I is the indicator function, and RWR_empirical_ and RWR_random_ refer to the RWR score assigned to the empirical PIN and the random networks, respectively. Only nodes with a p < 0.05 were selected. We called this subnetwork disease network.

### KO network and target gene prioritisation

To simulate the KOs on the upregulated network, we removed one of the DEG at a time from the initial PIN and the random networks and repeated the same procedure employed above. We generated 81 KO networks, and each KO network was similar to the empirical one, except that a node was missing. This manoeuvre permits mimicking the effect of a drug inhibitor on a protein. To prioritize the drug targets, we used i) Jensen-Shannon distance (Endres and Schindelin 2003):

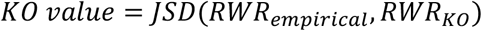

Where JSD refers to the Jensen-Shannon distance, *RWR*_*empirical*_ and *RWR*_*KO*_ represent the RWR vector obtained from the empirical and the KO networks, respectively—A more significant KO value indicates that the inhibition of protein causes a substantial effect on the empirical network; (ii) the RWR value of the empirical network; (iii) the betweenness centrality extracted from the disease network; and (iv) the DisGeneNet score to identify genes previously associated with CVDs such as atherosclerosis.

### Enrichment analysis of ranked ED gene list

We performed an over-representation analysis of the 81 KO ranked list of genes based on their JSD score and divided them into the top (41 genes) and bottom (40 genes) (Boyle et al., 2004). We performed the disease enrichment analysis using the R package enrichR for the DisGeneNet database (Chen et al., 2013). The enriched disease terms were ranked by their combined score, which is calculated by the c = log(p) * z, where *c* is the combined score, *p* is the Fisher exact test p-value, and *z* is the z-score for the deviation of expected rank (Chen et al., 2013).

### ChEMBL and OpenTarget drugs selection

Next, we used the OpenTargets (Ochoa et al. 2021) and ChEMBL (Willighagen et al. 2013) database to identify potent known inhibitors/antagonists for prioritised targets above. Each target had a list of active compounds with increasing values of Kd/Ki - or IC_50_ if Kd/Ki data was not available. The active compound with the lowest Kd/Ki/IC_50_ was then checked for assay data quality (assay type, cell type), physicochemical properties (Log P and molecular weight) (Zhu et al. 2013) and commercial availability (Sterling and Irwin 2015).

### Cell culture

HAECs were purchased from LONZA (Lonza, Walkersville, MA, USA). The cells were grown in medium EGM-2MV (LONZA), incubated at 37 ºC and 5% CO_2_. All experiments were performed between passages four and seven.

### Cytotoxicity assay of selected molecules

The 17 inhibitory or antagonistic molecules were tested for cytotoxicity using HAECs in the static culture at 10^−6^M and 10^−8^M. The higher dose (10^−6^M) was chosen to increase the chances of target inhibition, and the 100x lower dose (10^−8^ M) was chosen to reach concentrations closer to those presented in the circulation. We used the LIVE/DEAD® Viability/Cytotoxicity kit (Thermo Fisher Scientific) for this assay. HAECs at 100% confluence were seeded in 96-well plates (3.0 × 10^4^/well) and treated with the molecules at both concentrations. After 48 h, the medium was carefully removed, and the fluorescent probes were added to react with live (calcein-AM, 2 µM, Ex/Em 494/517) and dead (ethidium homodimer-1, 4 µM Ex/Em 528/617) cells, with a cell nucleus marker (Hoechst 8.1µM – Ex/Em 361/486) (Thermo Fisher Scientific) diluted in phosphate-buffered saline (PBS). The probes were incubated for 30 min at room temperature and then removed. Cells were fixed with 4% PFA for 15 minutes at room temperature. Finally, cells were maintained in PBS for image acquisition with the EVOS M7000 fluorescence microscope (Thermo Fisher Scientific). Nine pictures were taken per well at 20x magnification. We used HAECs stimulated with DMSO (at the same proportion of the highest dose 10^−6^ M of the molecules) as basal control. As a positive death control, cells were cultured in a starving medium for 24 h. The cell death index (ethidium and calcein index) was calculated by counting the number of positive-staining cells divided by the total number of Hoechst-positive cells multiplied by 100.

### High-content screening (HCS)

We evaluated the effect of the 17 drugs on ROS generation and induction of the inflammation protein ICAM-1. HAECs were plated in 96-well plates (3 × 10^4^ cells per well) for screening. We pre-treated the cells with 17 drugs diluted in a complete culture medium (EGM-2MV) at 10^−6^ M and 10^−8^ M for 2 h. Then, we added IL-1β (10 ng/mL, Peprotech) and OxPAPC (50 µg/mL, Hycult Biotech) to the medium. HAECs were cultured for 50 h (2 h pre-treatment with drugs + 48 h with drugs and stimulation with IL-1β and OxPAPC). We used as baseline controls HAECs at complete medium without DMSO, with DMSO (same proportion of drugs at 10^−6^ M), and with IL-1β and OxPAPC (without drugs). Additionally, we used PMA (10 µM, Sigma) and H_2_O_2_ (200 µM, Sigma) for 1 h before the end of the incubation period to stimulate ROS production. We used atorvastatin (5 µM, Sigma) to compare with the molecules tested. We followed the same approach, 2 h pre-treatment with atorvastatin (5 µM), followed by 48-h stimulation with IL-1β and OxPAPC.

After the end of the experiment, the cells were incubated with CM-H2DCFDA (5 µM, Thermo Fisher Scientific) and MitoSox (5 µM, Thermo Fisher Scientific) for 30 minutes at 37 ºC to evaluate ROS generation. The cells were fixed with 4% PFA for 15 minutes at room temperature. We followed the immunofluorescence protocol with permeabilisation for 1 h with Triton (0.1% in PBS), blocking with 5% bovine serum albumin in PBS for 1 h at room temperature, and incubating with the anti-ICAM1 antibody (Cell Signaling #62133/ 1:200), anti-E-selectin antibody (Abcam#ab18991/ 1:100) at 4 ºC overnight. After this incubation, we washed with PBS (2x). We followed the incubation with DAPI (1 mg/mL, Thermo Fisher Scientific) and fluorescent secondary antibodies (Alexa Fluor, Thermo Fisher Scientific, 1:500). The incubation of MitoSox and anti-E-selectin was performed only for EGLN3 inhibitor, after drug screening and analysis. Finally, we performed two more washes with PBS, and the cells were maintained in PBS for image acquisition.

### Image acquisition and analysis

We used an EVOS M7000 fluorescence microscope (Thermo Fisher Scientific). Images were obtained at 10x magnification, with nine images per well. After the acquisition, the raw images were analysed using CellProfiler (McQuin et al., 2018) software. The image analysis pipeline proceeded with identifying primary objects to identify the nucleus. Then, we identified and segmented the secondary objects for CM-H2DCFDA, MitoSox, ICAM-1, and E-selectin performed from DAPI markings. Next, we quantified the signal intensity for the identified secondary objects. We considered the median intensity for CM-H2DCFDA and MitoSox, the median edge intensity (EDGE) for ICAM-1, and cytoplasmatic staining for E-selectin. We considered the median intensity of each well. We also used the EDGE for ICAM-1 to calculate the eccentricity as the parameter to measure cell morphology. Parameters measured in addition to the intensity signal included cell area, length, and circularity using the ICAM-1 marker.

### Image data analysis

We performed the normalisation of the median intensities of CM-H2DCFDA and ICAM1 markers quantified using CellProfile (McQuin et al., 2018), with DMSO controls [normalised values = median intensity value/mean (median intensity DMSO control samples)] to normalize the median intensity of each well’s values within each plate. Then, to consider the plate-to-plate variation, we computed the z-score for all plates by subtracting the mean and dividing by the standard deviation of all samples. P-values were computed by multiple pairwise comparisons with the Wilcon test followed by BH correction, where we compared all conditions (DMSO, 10^−6^ M, and 10^−8^ M of the 17 drugs) vs HAECs treated with IL-1B&OxPAPC.

### Protein extraction, western blot, and analysis

Cells were lysed in RIPA buffer (150 mM NaCl, 0.5% sodium deoxycholate, 1% Triton X-100, 0.1% SDS, 50 mM Tris-HCl, pH 8) containing protease and phosphatase inhibitors (Sigma-Aldrich) and proteins quantified using a BCA protein assay kit (Pierce Biotechnology). Equivalent amounts of proteins were solubilised in sample buffer (0.5% SDS, 10% glycerol, 0.05% bromophenol blue, 50 mM dithiothreitol, 50 mM Tris, pH 6) and subjected to sodium dodecyl sulfate-polyacrylamide gel electrophoresis. After electrophoresis, the proteins were transferred to polyvinylidene fluoride membranes (Amersham Hybond). The membranes were first incubated in blocking solution (5% bovine serum albumin and 0.1% Tween 20 in tris-buffered saline, pH 7.4) for 1 h at room temperature, followed by incubation for 16–18 h in primary antibody against HIF-1a (Cell Signalling #36169/1:500) diluted in blocking solution at 4°C. Membranes were also incubated with primary antibodies specific for GAPDH (Abcam#ab22555, 1:2000) as an internal control (normaliser). After incubation for 1 h at room temperature with a secondary antibody conjugated to horseradish peroxidase, the bound primary antibody was detected using a chemiluminescent image analyser (ImageQuant LAS 4000 mini), and images were quantified by densitometry using ImageJ (Schneider, Rasband, and Eliceiri, 2012).

### qPCR and analysis

ECs were lysed with 1 mL of TRIzol Reagent (Invitrogen), and the total RNA was extracted according to the manufacturer’s instructions. After extraction and purification, RNA was reverse transcribed using SuperScript IV Reverse Transcriptase (Thermo Fisher Scientific). Real-time PCR was performed using SYBR-Green (Roche) with the QuantiStudio 12 K Flex system (Applied Biosystems). We used GAPDH Ct values to normalise ΔCt. The variation in gene expression between samples was calculated using the ΔΔCt method.

The following primers were used (5′ to 3′):

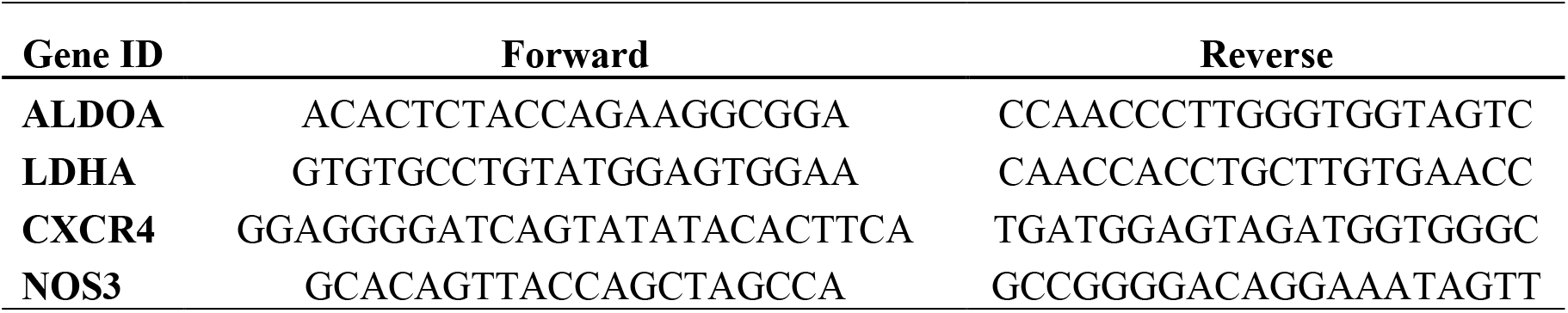

### Statistical analysis

For qPCR and western blot, we used Student’s t-test or one-way analysis of variance, followed by the Benjamini–Hochberg (BH) post hoc test. We analysed variance using Kruskal-Wallis ranks for nonparametric values, followed by the Wilcoxon test for multiple comparisons. Differences were significant when p < 0.05. For all statistical analyses, we used R software (version 4.1.0). All the experiments were performed at least two times (one biological replicate for each time) with 4 to 5 technical replicates. Outliers were removed based on the z-score methods with an absolute value of 3.

## Supplemental material

**Supplementary Fig1.**
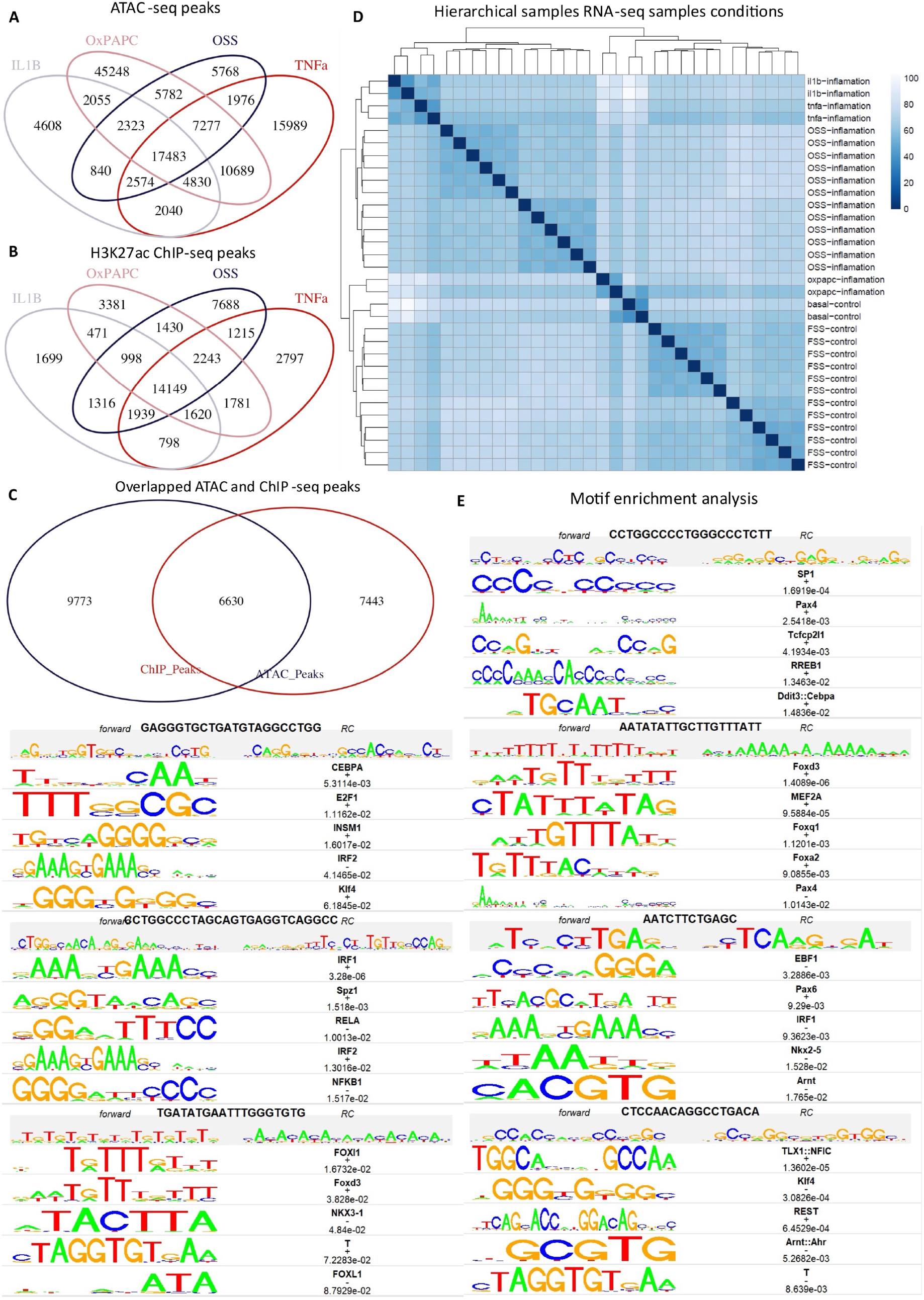
Surrogates of CV risk factors OSS, IL-1β, TNF-α, and OxPAPC modulate common regulatory regions on the genome. **(A)** Venn diagram of ATAC-seq peaks of all conditions. **(B)** Venn diagram of ChIP-seq peaks of all conditions. **(C)** Venn diagram of the overlapped peaks of ATAC-seq and ChIP-seq. **(D)** The hierarchical clustering profile reveals complex relationships between all samples under IL-1β, TNF-α, OSS, OxPAPC, basal control, and FSS conditions. The heat map of all samples using Euclidean distance as a measured parameter. **(E)** Enriched motifs in the centre of enhancer-like regions are shown, including the transcription factor family and motif sequence at enrichment log P-values. Enrichment was calculated from 200 bp sequences centred on chromatin accessibility.

**Supplementary Fig 2.**
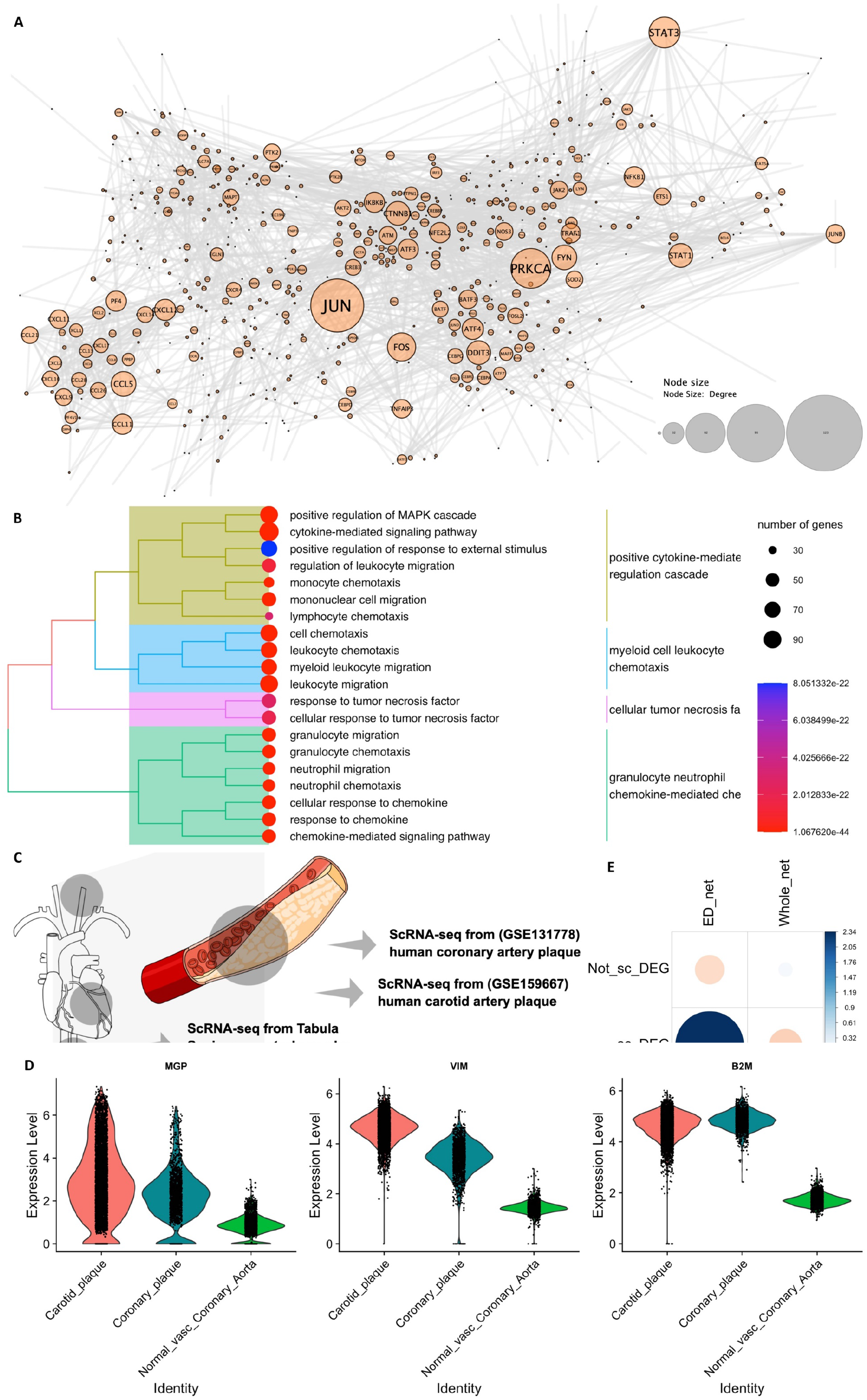
TF-DEGs seed nodes. (**A**) Protein-interaction network, ED disease network generated from the RWR algorithm. The network comprises 838 nodes and 1,991 edges. The size of the node represents the degree, i.e., the number of interacting partners (**B**) GO enrichment analysis for BP for all 838 nodes, followed by redundancy analysis to cluster the terms with high semantic similarity (adjusted p ≤ 0.05). **(C)** Schematic figure showing the scRNA-seq public data used (Methods) to validate the ED disease network. The data comprised two atherosclerosis lesion datasets and one Tabula sapiens control dataset. The cartoons were created using the Mind the Graph platform (www.mindthegraph.com). **(D)** Violin plot of the three top-upregulated genes in the ECs from atherosclerotic sites compared to the normal vasculature. **(E)** Hypergeometric test p = 0.008. Positive residuals are in blue, and negative residuals are in red.

**Supplementary Fig 3.**
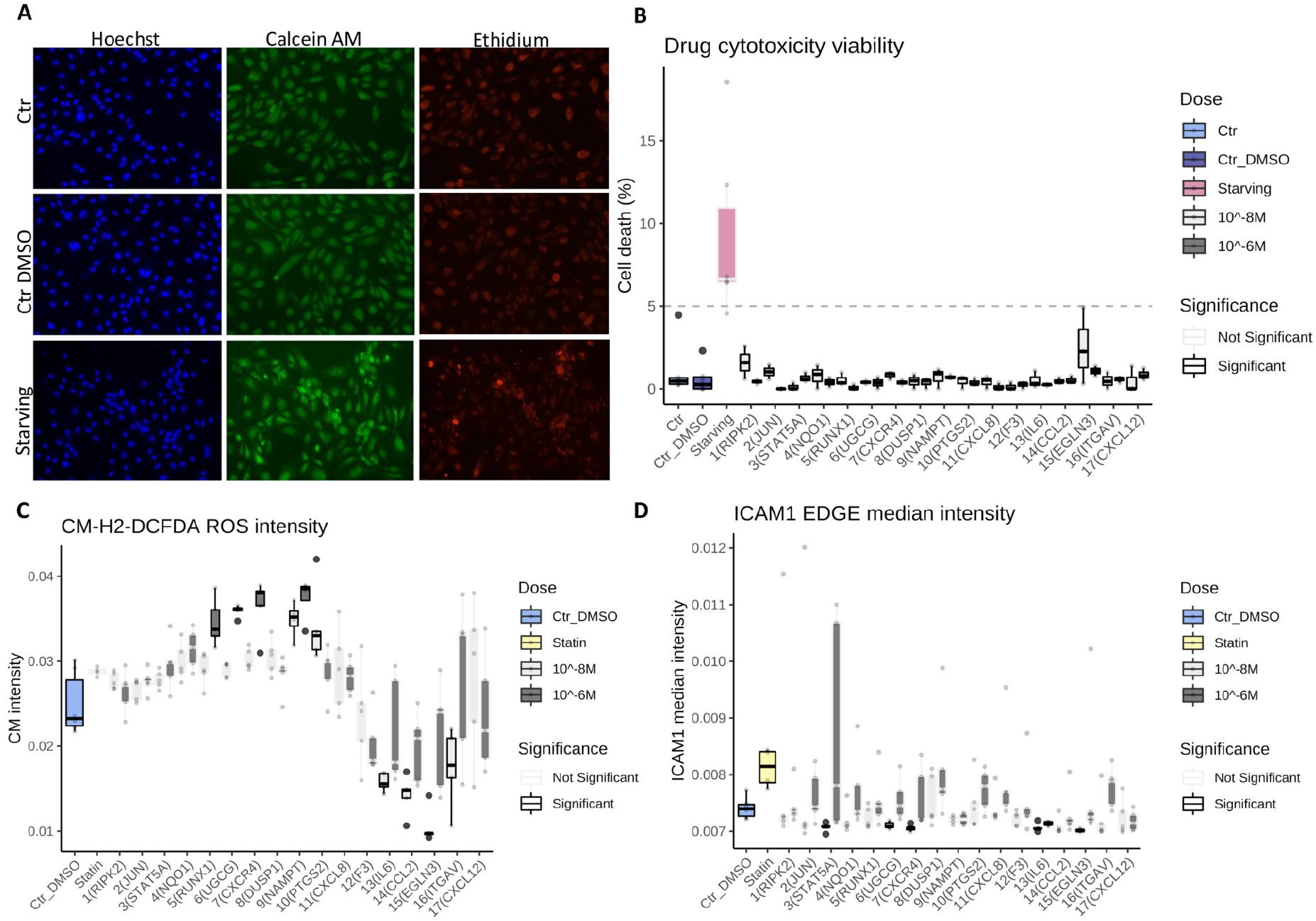
Cytotoxicity test of the 17 drugs tested at 10^−6^ M and 10^−8^ M. **(A)** Representative fluorescent images of the viability assay. We used starving treatment for 24 h as a cell death positive control. The nuclei staining is in blue (Hoechst), live cells are stained in green (calcein), while the nuclei of dead cells are marked with ethidium, red staining. **(B)** The plot shows all 17 drugs tested at 10^−8^M and 10^−6^M. The percentage of cell death was calculated by the number of dead/live cells relative to the number of cells (nuclei). **(C)** The plot shows the intensity of the signals for the probe CM-H2-DCFDA – (green), a ROS indicator **(D)** and the intensity of ICAM1 staining (antibody anti-ICAM1, red). P-values were computed by multiple pairwise comparisons with the Wilcon test followed by BH correction, where all conditions were compared to starving (cell viability assay) or DMSO (Figure C, D). Data are presented as mean ± standard deviation. The experiment has one biological replicate with 3 technical replicates.

**Supplementary Figure 4.**
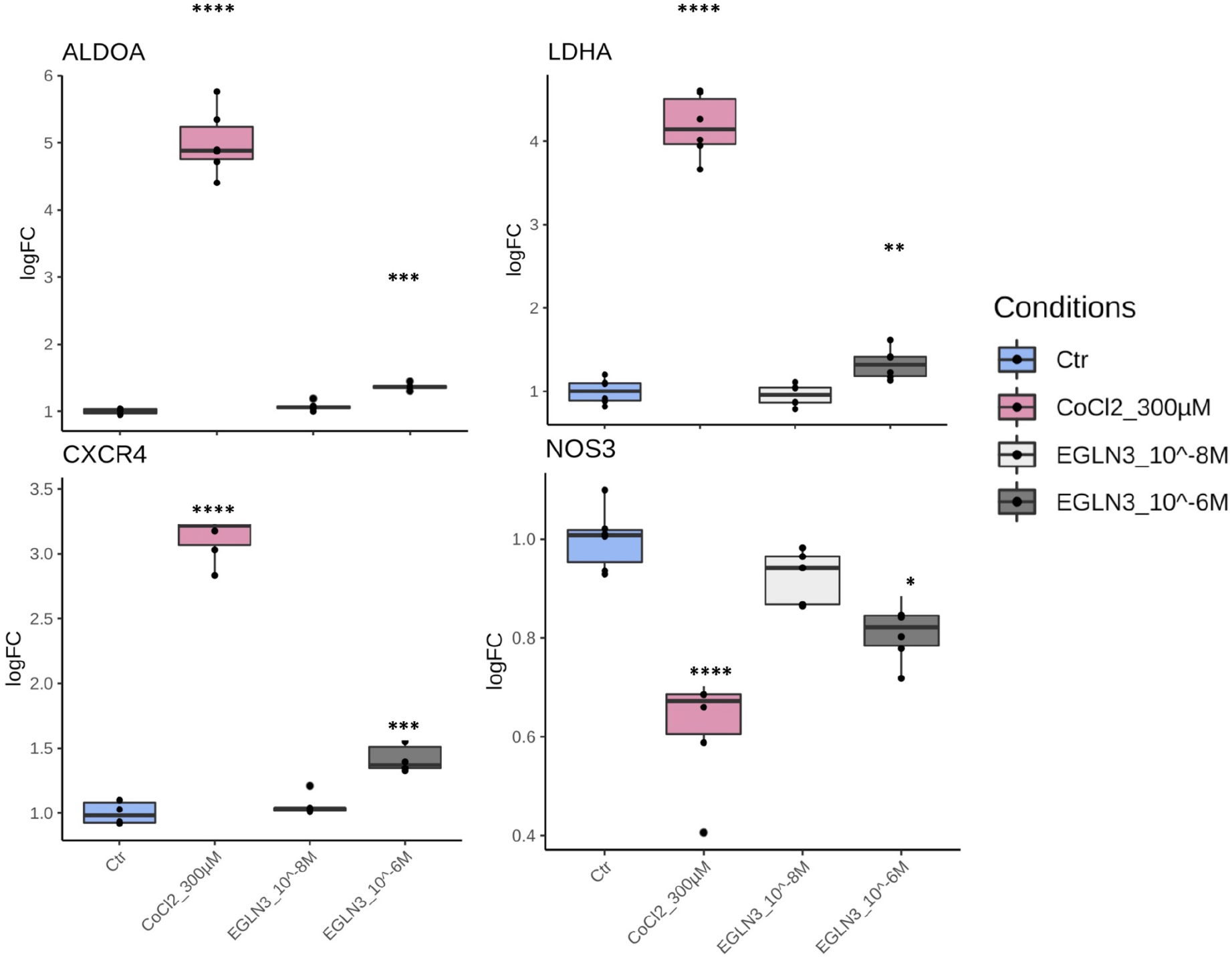
Molidustat HIF-1α downstream gene target modulation. HAEC were treated with Molidustat at 10^−8^ M and 10^−6^ M, or cobalt chloride (CoCl_2_, 300 μM) for 48 h and gene expression of aldolase, fructose-bisphosphate A (ALDOA), lactate dehydrogenase A (LDHA), C-X-C motif chemokine receptor 4 (CXCR4) and nitric oxide synthase 3 (NOS3) were evaluated using qPCR. Data are presented as mean ± EPM. The experiment has 4 biological replicates with 2 technical replicates of each condition.

**Extended blot Fig 4F.**
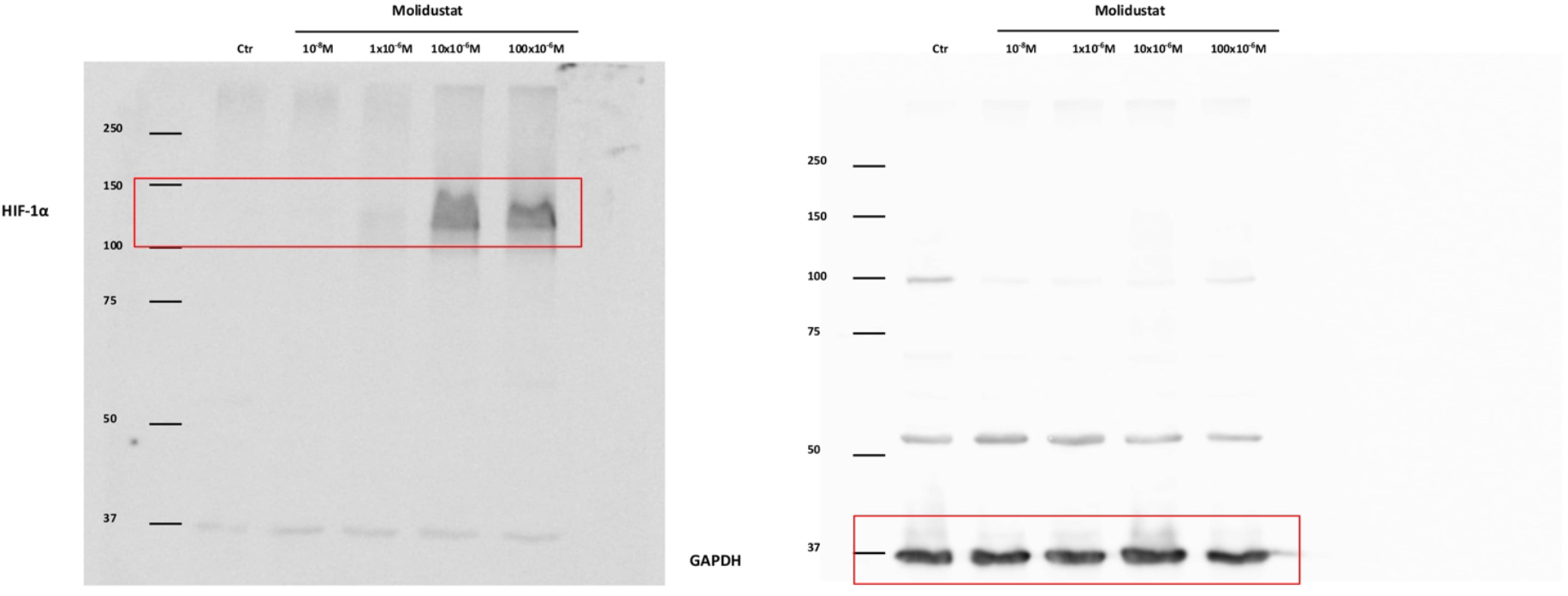
HAEC were treated with Molidustat at 10^−8^ M to 100 × 10^−6^ M for 48 h. HIF-1α protein expression was determined by western blot. GAPDH served as a housekeeping protein.

**Supplementary Table 5.**
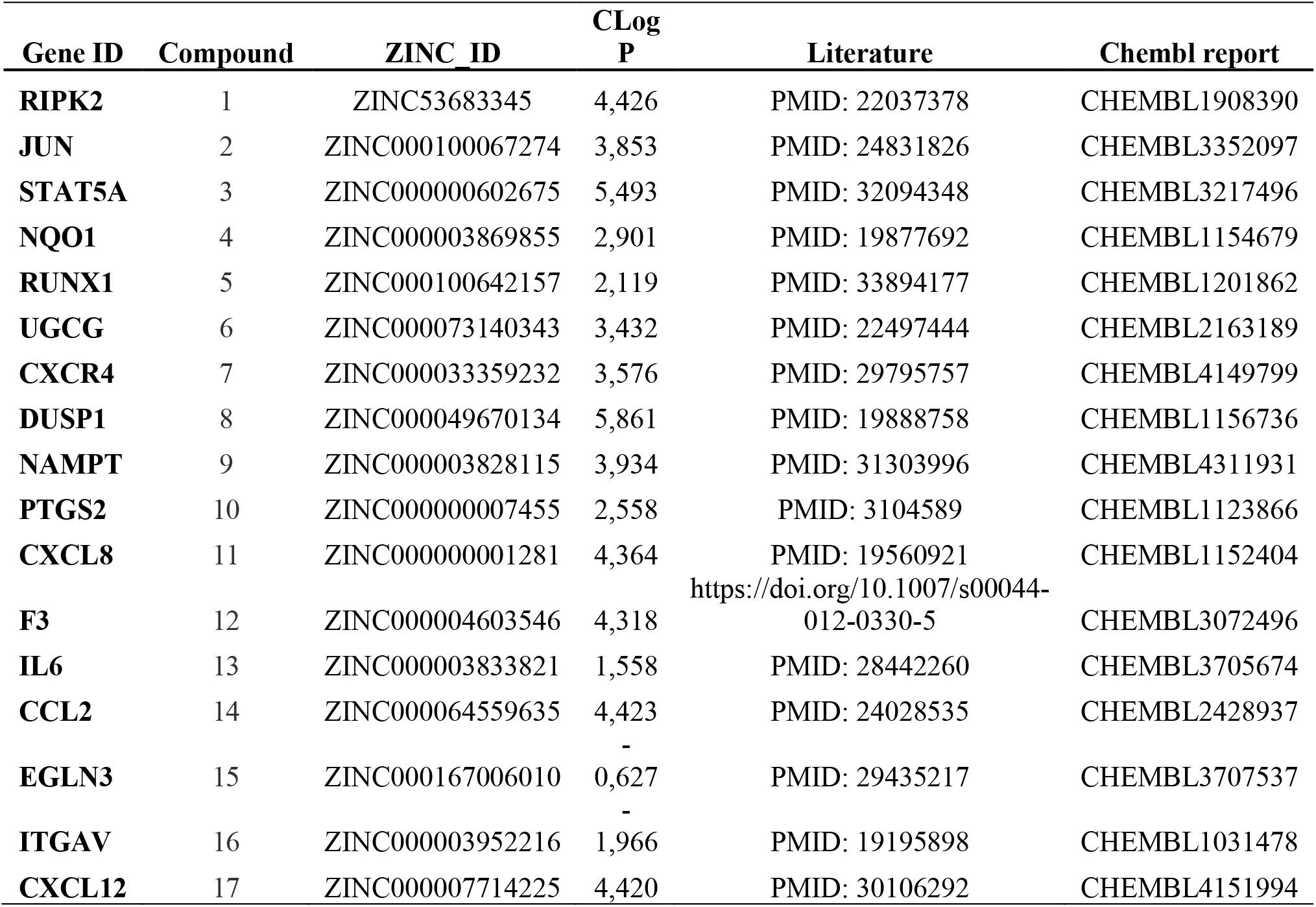
Description of the seventeen small molecules that target the selected genes candidates

## References

Abe, Hajime, Hiroaki Semba, and Norihiko Takeda. 2017. “The Roles of Hypoxia Signaling in the Pathogenesis Of.” 1:884–94.

Alsaigh, Tom, Doug Evans, David Frankel, and Ali Torkamani. 2020. “Decoding the Transcriptome of Atherosclerotic Plaque at Single-Cell Resolution.” doi: 10.1101/2020.03.03.968123.

Appelhoffl, Rebecca J., Ya Min Tian, Raju R. Raval, Helen Turley, Adrian L. Harris, Christopher W. Pugh, Peter J. Ratcliffe, and Jonathan M. Gleadle. 2004. “Differential Function of the Prolyl Hydroxylases PHD1, PHD2, and PHD3 in the Regulation of Hypoxia-Inducible Factor.” Journal of Biological Chemistry. doi: 10.1074/jbc.M406026200.

Beck, Hartmut, Mario Jeske, Kai Thede, Friederike Stoll, Ingo Flamme, Metin Akbaba, Jens Kerim Ergüden, Gunter Karig, Jörg Keldenich, Felix Oehme, Hans Christian Militzer, Ingo v. Hartung, and Uwe Thuss. 2018. “Discovery of Molidustat (BAY 85-3934): A Small-Molecule Oral HIF-Prolyl Hydroxylase (HIF-PH) Inhibitor for the Treatment of Renal Anemia.” ChemMedChem 13(10):988–1003. doi: 10.1002/cmdc.201700783.

Boulanger, Chantal M. 2016. “Endothelium.” Arteriosclerosis, Thrombosis, and Vascular Biology. doi: 10.1161/ATVBAHA.116.306940.

Castro-Mondragon, Jaime A., Rafael Riudavets-Puig, Ieva Rauluseviciute, Roza Berhanu Lemma, Laura Turchi, Romain Blanc-Mathieu, Jeremy Lucas, Paul Boddie, Aziz Khan, Nicolás Manosalva Perez, Oriol Fornes, Tiffany Y. Leung, Alejandro Aguirre, Fayrouz Hammal, Daniel Schmelter, Damir Baranasic, Benoit Ballester, Albin Sandelin, Boris Lenhard, Klaas Vandepoele, Wyeth W. Wasserman, François Parcy, and Anthony Mathelier. 2022. “JASPAR 2022: The 9th Release of the Open-Access Database of Transcription Factor Binding Profiles.” Nucleic Acids Research 50(D1):D165–73. doi: 10.1093/nar/gkab1113.

Cava, Claudia, and Isabella Castiglioni. 2019. “In Silico Perturbation of Drug Targets in Pan-Cancer Analysis Combining Multiple Networks and Pathways.” Gene 698(February):100–106. doi: 10.1016/j.gene.2019.02.064.

Chen, Edward Y., Christopher M. Tan, Yan Kou, Qiaonan Duan, Zichen Wang, Gabriela Vaz Meirelles, Neil R. Clark, and Avi Ma’ayan. 2013. Enrichr: Interactive and Collaborative HTML5 Gene List Enrichment Analysis Tool.

Chiu, Jeng Jiann, Pei Ling Lee, Cheng Nan Chen, Chih I. Lee, Shun Fu Chang, Li Jing Chen, Sheng Chieh Lien, Ya Chen Ko, Shunichi Usami, and Shu Chien. 2004. “Shear Stress Increases ICAM-1 and Decreases VCAM-1 and E-Selectin Expressions Induced by Tumor Necrosis Factor-α in Endothelial Cells.” Arteriosclerosis, Thrombosis, and Vascular Biology 24(1):73–79. doi: 10.1161/01.ATV.0000106321.63667.24.

Cyr, Anthony R., Lauren v. Huckaby, Sruti S. Shiva, and Brian S. Zuckerbraun. 2020. “Nitric Oxide and Endothelial Dysfunction.” Critical Care Clinics.

Daiber, Andreas, and Stefan Chlopicki. 2020. “Revisiting Pharmacology of Oxidative Stress and Endothelial Dysfunction in Cardiovascular Disease: Evidence for Redox-Based Therapies.” Free Radical Biology and Medicine 157:15–37.

Daiber, Andreas, Sebastian Steven, Alina Weber, Vladimir v. Shuvaev, Vladimir R. Muzykantov, Ismail Laher, Huige Li, Santiago Lamas, and Thomas Münzel. 2017. “Targeting Vascular (Endothelial) Dysfunction.” British Journal of Pharmacology 174(12):1591–1619. doi: 10.1111/bph.13517.

Davidich, Maria I., and Stefan Bornholdt. 2013. “Boolean Network Model Predicts Knockout Mutant Phenotypes of Fission Yeast.” PLoS ONE 8(9). doi: 10.1371/journal.pone.0071786.

Deanfield, John E., Julian P. Halcox, and Ton J. Rabelink. 2007. “Endothelial Function and Dysfunction: Testing and Clinical Relevance.” Circulation 115(10):1285–95. doi: 10.1161/CIRCULATIONAHA.106.652859.

Dorel, Mathurin, Bertram Klinger, Torsten Gross, Anja Sieber, Anirudh Prahallad, Evert Bosdriesz, Lodewyk F. A. Wessels, and Nils Blüthgen. 2018. “Modelling Signalling Networks from Perturbation Data.” Bioinformatics 34(23):4079–86. doi: 10.1093/bioinformatics/bty473.

Endres, Dominik M., and Johannes E. Schindelin. 2003. “A New Metric for Probability Distributions.” IEEE Transactions on Information Theory 49(7):1858–60.

Fang, Yun, David Wu, and Konstantin G. Birukov. 2019. “Mechanosensing and Mechanoregulation of Endothelial Cell Functions.” Comprehensive Physiology 9(2):873–904. doi: 10.1002/cphy.c180020.

Feng, Y., Z. R. Cai, Y. Tang, G. Hu, J. Lu, D. He, and S. Wang. 2014. “TLR4/NF-?B Signaling Pathway-Mediated and OxLDL-Induced up-Regulation of LOX-1, MCP-1, and VCAM-1 Expressions in Human Umbilical Vein Endothelial Cells.” Genetics and Molecular Research 13(1):680–95.

Fisker Hag, Anne Mette, Sune Folke Pedersen, and Andreas Kjaer. 2008. “Gene Expression of LOX-1, VCAM-1, and ICAM-1 in Pre-Atherosclerotic Mice.” Biochemical and Biophysical Research Communications 377(2):689–93. doi: 10.1016/j.bbrc.2008.10.037.

Gencer, Selin, Bryce R. Evans, Emiel P. C. van der Vorst, Yvonne Döring, and Christian Weber. 2021. “Inflammatory Chemokines in Atherosclerosis.” 1–26.

Godo, Shigeo, and Hiroaki Shimokawa. 2017. “Endothelial Functions.” Arteriosclerosis, Thrombosis, and Vascular Biology 37(9):e108–14. doi: 10.1161/ATVBAHA.117.309813.

Gokce, Noyan, John F. Keaney, Liza M. Hunter, Michael T. Watkins, James O. Menzoian, and Joseph A. Vita. 2002. “Risk Stratification for Postoperative Cardiovascular Events via Noninvasive Assessment of Endothelial Function: A Prospective Study.” Circulation 105(13):1567–72. doi: 10.1161/01.CIR.0000012543.55874.47.

Han, Heonjong, Jae Won Cho, Sangyoung Lee, Ayoung Yun, Hyojin Kim, Dasom Bae, Sunmo Yang, Chan Yeong Kim, Muyoung Lee, Eunbeen Kim, Sungho Lee, Byunghee Kang, Dabin Jeong, Yaeji Kim, Hyeon Nae Jeon, Haein Jung, Sunhwee Nam, Michael Chung, Jong Hoon Kim, and Insuk Lee. 2018. “TRRUST v2: An Expanded Reference Database of Human and Mouse Transcriptional Regulatory Interactions.” Nucleic Acids Research. doi: 10.1093/nar/gkx1013.

Hennigs, Jan K., Christiane Matuszcak, Martin Trepel, and Jakob Körbelin. 2021. “Vascular Endothelial Cells: Heterogeneity and Targeting Approaches.” Cells 10(10).

Herrmann, Stefan-Martin, Carl Whatling, Eva Brand, Viviane Nicaud, Jérôme Gariepy, Alain Simon, Alun Evans, Jean-Bernard Ruidavets, Dominique Arveiler, Gerald Luc, Laurence Tiret, Adriano Henney, and François Cambien. 2000. Polymorphisms of the Human Matrix Gla Protein (MGP) Gene, Vascular Calcification, and Myocardial Infarction.

Hogan, Nicholas T., Michael B. Whalen, Lindsey K. Stolze, Nizar K. Hadeli, Michael T. Lam, James R. Springstead, Christopher K. Glass, and Casey E. Romanoski. 2017. “Transcriptional Networks Specifying Homeostatic and Inflammatory Programs of Gene Expression in Human Aortic Endothelial Cells.” (Cvd):1–28. doi: 10.7554/eLife.22536.

Hornbeck, Peter v., Bin Zhang, Beth Murray, Jon M. Kornhauser, Vaughan Latham, and Elzbieta Skrzypek. 2015. “PhosphoSitePlus, 2014: Mutations, PTMs and Recalibrations.” Nucleic Acids Research. doi: 10.1093/nar/gku1267.

Huynh, Diem Thi Ngoc, and Kyung Sun Heo. 2019. “Therapeutic Targets for Endothelial Dysfunction in Vascular Diseases.” Archives of Pharmacal Research 42(10):848–61.

Inflammasome, The, and Peter Libby. 2021. “Targeting Inflammatory Pathways in Cardiovascular Disease:”

Joharapurkar, Amit A., Vrajesh B. Pandya, Vishal J. Patel, Ranjit C. Desai, and Mukul R. Jain. 2018. “Prolyl Hydroxylase Inhibitors: A Breakthrough in the Therapy of Anemia Associated with Chronic Diseases.” Journal of Medicinal Chemistry 61(16):6964–82.

Jones, Robert C., Jim Karkanias, Mark A. Krasnow, Angela Oliveira Pisco, Stephen R. Quake, Julia Salzman, Nir Yosef, Bryan Bulthaup, Phillip Brown, William Harper, Marisa Hemenez, Ravikumar Ponnusamy, Ahmad Salehi, Bhavani A. Sanagavarapu, Eileen Spallino, Ksenia A. Aaron, Waldo Concepcion, James M. Gardner, Burnett Kelly, Nikole Neidlinger, Zifa Wang, Sheela Crasta, Saroja Kolluru, Maurizio Morri, Serena Y. Tan, Kyle J. Travaglini, Chenling Xu, Marcela Alcántara-Hernández, Nicole Almanzar, Jane Antony, Benjamin Beyersdorf, Deviana Burhan, Kruti Calcuttawala, Matthew M. Carter, Charles K. F. Chan, Charles A. Chang, Stephen Chang, Alex Colville, Rebecca N. Culver, Ivana Cvijović, Gaetano D’Amato, Camille Ezran, Francisco X. Galdos, Astrid Gillich, William R. Goodyer, Yan Hang, Alyssa Hayashi, Sahar Houshdaran, Xianxi Huang, Juan C. Irwin, So Ri Jang, Julia Vallve Juanico, Aaron M. Kershner, Soochi Kim, Bernhard Kiss, William Kong, Maya E. Kumar, Angera H. Kuo, Rebecca Leylek, Baoxiang Li, Gabriel B. Loeb, Wan Jin Lu, Sruthi Mantri, Maxim Markovic, Patrick L. McAlpine, Antoine de Morree, Karim Mrouj, Shravani Mukherjee, Tyler Muser, Patrick Neuhöfer, Thi D. Nguyen, Kimberly Perez, Ragini Phansalkar, Nazan Puluca, Zhen Qi, Poorvi Rao, Hayley Raquer-McKay, Nicholas Schaum, Bronwyn Scott, Bobak Seddighzadeh, Joe Segal, Sushmita Sen, Shaheen Sikandar, Sean P. Spencer, Lea C. Steffes, Varun R. Subramaniam, Aditi Swarup, Michael Swift, Will van Treuren, Emily Trimm, Stefan Veizades, Sivakamasundari Vijayakumar, Kim Chi Vo, Sevahn K. Vorperian, Wanxin Wang, Hannah N. W. Weinstein, Juliane Winkler, Timothy T. H. Wu, Jamie Xie, Andrea R. Yung, Yue Zhang, Angela M. Detweiler, Honey Mekonen, Norma F. Neff, Rene v. Sit, Michelle Tan, Jia Yan, Gregory R. Bean, Vivek Charu, Erna Forgó, Brock A. Martin, Michael G. Ozawa, Oscar Silva, Angus Toland, Venkata N. P. Vemuri, Shaked Afik, Kyle Awayan, Olga Borisovna Botvinnik, Ashley Byrne, Michelle Chen, Roozbeh Dehghannasiri, Angela M. Detweiler, Adam Gayoso, Alejandro A. Granados, Qiqing Li, Gita Mahmoudabadi, Aaron McGeever, Julia Eve Olivieri, Madeline Park, Neha Ravikumar, Geoff Stanley, Michelle Tan, Weilun Tan, Alexander J. Tarashansky, Rohan Vanheusden, Sevahn K. Vorperian, Peter Wang, Sheng Wang, Galen Xing, Les Dethlefsen, Po Yi Ho, Shixuan Liu, Jonathan S. Maltzman, Ross J. Metzger, Kimberly Perez, Hayley Raquer-McKay, Koki Sasagawa, Rahul Sinha, Hanbing Song, Aditi Swarup, Sivakamasundari Vijayakumar, Bruce Wang, Andrea R. Yung, Steven E. Artandi, Philip A. Beachy, Michael F. Clarke, Linda C. Giudice, Franklin W. Huang, Kerwyn Casey Huang, Juliana Idoyaga, Seung K. Kim, Christin S. Kuo, Patricia Nguyen, Thomas A. Rando, Kristy Red-Horse, Jeremy Reiter, David A. Relman, Justin L. Sonnenburg, Albert Wu, Sean M. Wu, and Tony Wyss-Coray. 2022. “The Tabula Sapiens: A Multiple-Organ, Single-Cell Transcriptomic Atlas of Humans.” Science 376(6594). doi: 10.1126/science.abl4896.

Kerrien, Samuel, Bruno Aranda, Lionel Breuza, Alan Bridge, Fiona Broackes-Carter, Carol Chen, Margaret Duesbury, Marine Dumousseau, Marc Feuermann, Ursula Hinz, Christine Jandrasits, Rafael C. Jimenez, Jyoti Khadake, Usha Mahadevan, Patrick Masson, Ivo Pedruzzi, Eric Pfeiffenberger, Pablo Porras, Arathi Raghunath, Bernd Roechert, Sandra Orchard, and Henning Hermjakob. 2012. “The IntAct Molecular Interaction Database in 2012.” Nucleic Acids Research. doi: 10.1093/nar/gkr1088.

Kim, Tae Hoon, and Job Dekker. 2018. “ChIP-Seq.” Cold Spring Harbor Protocols. doi: 10.1101/pdb.prot082644.

Krause, Matthew D., Ru-ting Huang, David Wu, Tzu-pin Shentu, Devin L. Harrison, and Michael B. Whalen. 2018. “Genetic Variant at Coronary Artery Disease and Ischemic Stroke Locus 1p32. 2 Regulates Endothelial Responses to Hemodynamics.” (6). doi: 10.1073/pnas.1810568115.

Langmead, Ben, and Steven L. Salzberg. 2012. “Fast Gapped-Read Alignment with Bowtie 2.” Nature Methods 9(4):357–59. doi: 10.1038/nmeth.1923.

Lawrence, Michael, Robert Gentleman, and Vincent Carey. 2009. “Rtracklayer: An R Package for Interfacing with Genome Browsers.” Bioinformatics 25(14):1841–42. doi: 10.1093/bioinformatics/btp328.

Lawrence, Michael, Wolfgang Huber, Hervé Pagès, Patrick Aboyoun, Marc Carlson, Robert Gentleman, Martin T. Morgan, and Vincent J. Carey. 2013. “Software for Computing and Annotating Genomic Ranges.” PLoS Computational Biology 9(8). doi: 10.1371/journal.pcbi.1003118.

Ley, Klaus, Carlo Laudanna, Myron I. Cybulsky, and Sussan Nourshargh. 2007. “Getting to the Site of Inflammation: The Leukocyte Adhesion Cascade Updated.” Nature Reviews Immunology.

Lin, Sean, Kenji Schorpp, Ina Rothenaigner, and Kamyar Hadian. 2020. “Image-Based High-Content Screening in Drug Discovery.” Drug Discovery Today 25(8):1348–61.

Liu, Weiwei, Yanlin Yin, Zihui Zhou, Min He, and Yalei Dai. 2014. “OxLDL-Induced IL-1beta Secretion Promoting Foam Cells Formation Was Mainly via CD36 Mediated ROS Production Leading to NLRP3 Inflammasome Activation.” Inflammation Research. doi: 10.1007/s00011-013-0667-3.

Liu, Ziqing, Dana L. Ruter, Kaitlyn Quigley, Natalie T. Tanke, Yuchao Jiang, and Victoria L. Bautch. 2021. “Single-Cell RNA Sequencing Reveals Endothelial Cell Transcriptome Heterogeneity Under Homeostatic Laminar Flow.” Arteriosclerosis, Thrombosis, and Vascular Biology (October):2575–84. doi: 10.1161/atvbaha.121.316797.

Love, Michael I., Wolfgang Huber, and Simon Anders. 2014. “Moderated Estimation of Fold Change and Dispersion for RNA-Seq Data with DESeq2.” Genome Biology 15(12). doi: 10.1186/s13059-014-0550-8.

Luo, Weibo, Hongxia Hu, Ryan Chang, Jun Zhong, Matthew Knabel, Robert O’Meally, Robert N. Cole, Akhilesh Pandey, and Gregg L. Semenza. 2011. “Pyruvate Kinase M2 Is a PHD3-Stimulated Coactivator for Hypoxia-Inducible Factor 1.” Cell. doi: 10.1016/j.cell.2011.03.054.

Lüscher, T. F., and M. Barton. 1997. “Biology of the Endothelium.” Clinical Cardiology. doi: 9422846.

Matsuzawa, Yasushi, Seigo Sugiyama, Hitoshi Sumida, Koichi Sugamura, Toshimitsu Nozaki, Keisuke Ohba, Junichi Matsubara, Hirofumi Kurokawa, Koichiro Fujisue, Masaaki Konishi, Eiichi Akiyama, Hiroyuki Suzuki, Yasuhiro Nagayoshi, Megumi Yamamuro, Kenji Sakamoto, Satomi Iwashita, Hideaki Jinnouchi, Masataka Taguri, Satoshi Morita, Kunihiko Matsui, Kazuo Kimura, Satoshi Umemura, and Hisao Ogawa. 2013. “Peripheral Endothelial Function and Cardiovascular Events in High-Risk Patients.” Journal of the American Heart Association 2(6). doi: 10.1161/JAHA.113.000426.

Mattiazzi Usaj, Mojca, Erin B. Styles, Adrian J. Verster, Helena Friesen, Charles Boone, and Brenda J. Andrews. 2016. “High-Content Screening for Quantitative Cell Biology.” Trends in Cell Biology 26(8):598–611. doi: 10.1016/j.tcb.2016.03.008.

McQuin, Claire, Allen Goodman, Vasiliy Chernyshev, Lee Kamentsky, Beth A. Cimini, Kyle W. Karhohs, Minh Doan, Liya Ding, Susanne M. Rafelski, Derek Thirstrup, Winfried Wiegraebe, Shantanu Singh, Tim Becker, Juan C. Caicedo, and Anne E. Carpenter. 2018. “CellProfiler 3.0: Next-Generation Image Processing for Biology.” PLoS Biology. doi: 10.1371/journal.pbio.2005970.

Mudau, Mashudu, Amanda Genis, Amanda Lochner, and Hans Strijdom. 2012. “Endothelial Dysfunction: The Early Predictor of Atherosclerosis.” Cardiovascular Journal of Africa 23(4):222–31. doi: 10.5830/CVJA-2011-068.

Mussbacher, Marion, Manuel Salzmann, Christine Brostjan, Bastian Hoesel, Christian Schoergenhofer, Hannes Datler, Philipp Hohensinner, José Basílio, Peter Petzelbauer, Alice Assinger, and Johannes A. Schmid. 2019. “Cell Type Specific Roles of Nf-Kb Linking Inflamation and Thrombosis.” Frontiers in Immunology 10(FEB).

Neckář, Jan, Anna Hsu, Md. Abdul Hye Khan, Garrett J. Gross, Kasem Nithipatikom, Michaela Cyprová, Daniel Benák, Markéta Hlaváčková, Dita Sotáková-Kašparová, John R. Falck, David Sedmera, František Kolář, and John D. Imig. 2018. “Infarct Size-Limiting Effect of Epoxyeicosatrienoic Acid Analog EET-B Is Mediated by Hypoxia-Inducible Factor-1α via Downregulation of Prolyl Hydroxylase 3.” American Journal of Physiology-Heart and Circulatory Physiology. doi: 10.1152/ajpheart.00726.2017.

Ochoa, David, Andrew Hercules, Miguel Carmona, Daniel Suveges, Asier Gonzalez-Uriarte, Cinzia Malangone, Alfredo Miranda, Luca Fumis, Denise Carvalho-Silva, Michaela Spitzer, Jarrod Baker, Javier Ferrer, Arwa Raies, Olesya Razuvayevskaya, Adam Faulconbridge, Eirini Petsalaki, Prudence Mutowo, Sandra MacHlitt-Northen, Gareth Peat, Elaine McAuley, Chuang Kee Ong, Edward Mountjoy, Maya Ghoussaini, Andrea Pierleoni, Eliseo Papa, Miguel Pignatelli, Gautier Koscielny, Mohd Karim, Jeremy Schwartzentruber, David G. Hulcoop, Ian Dunham, and Ellen M. McDonagh. 2021. “Open Targets Platform: Supporting Systematic Drug-Target Identification and Prioritisation.” Nucleic Acids Research 49(D1):D1302–10. doi: 10.1093/nar/gkaa1027.

Oesterle, Adam, Ulrich Laufs, and James K. Liao. 2017. “Pleiotropic Effects of Statins on the Cardiovascular System.” Circulation Research.

Orchard, Sandra, Mais Ammari, Bruno Aranda, Lionel Breuza, Leonardo Briganti, Fiona Broackes-Carter, Nancy H. Campbell, Gayatri Chavali, Carol Chen, Noemi Del-Toro, Margaret Duesbury, Marine Dumousseau, Eugenia Galeota, Ursula Hinz, Marta Iannuccelli, Sruthi Jagannathan, Rafael Jimenez, Jyoti Khadake, Astrid Lagreid, Luana Licata, Ruth C. Lovering, Birgit Meldal, Anna N. Melidoni, Mila Milagros, Daniele Peluso, Livia Perfetto, Pablo Porras, Arathi Raghunath, Sylvie Ricard-Blum, Bernd Roechert, Andre Stutz, Michael Tognolli, Kim van Roey, Gianni Cesareni, and Henning Hermjakob. 2014. “The MIntAct Project - IntAct as a Common Curation Platform for 11 Molecular Interaction Databases.” Nucleic Acids Research 42(D1). doi: 10.1093/nar/gkt1115.

Peiffer, Veronique, Spencer J. Sherwin, and Peter D. Weinberg. 2013. “Does Low and Oscillatory Wall Shear Stress Correlate Spatially with Early Atherosclerosis? A Systematic Review.” Cardiovascular Research.

Pinheiro-de-Sousa, Iguaracy, Miriam H. Fonseca-Alaniz, Samantha K. Teixeira, Mariliza v. Rodrigues, and Jose E. Krieger. 2022. “Uncovering Emergent Phenotypes in Endothelial Cells by Clustering of Surrogates of Cardiovascular Risk Factors.” Scientific Reports 12(1). doi: 10.1038/s41598-022-05404-7.

Premer, Courtney, Anthony J. Kanelidis, Joshua M. Hare, and Ivonne Hernandez Schulman. 2019. “Rethinking Endothelial Dysfunction as a Crucial Target in Fighting Heart Failure.” Mayo Clinic Proceedings: Innovations, Quality & Outcomes 3(1):1–13. doi: 10.1016/j.mayocpiqo.2018.12.006.

Putri, Givanna H., Simon Anders, Paul Theodor Pyl, John E. Pimanda, and Fabio Zanini. 2022. “Analysing High-Throughput Sequencing Data in Python with HTSeq 2.0.” Bioinformatics 38(10):2943–45. doi: 10.1093/bioinformatics/btac166.

Rahtu-Korpela, Lea, Jenni Määttä, Elitsa Y. Dimova, Sohvi Hörkkö, Helena Gylling, Gail Walkinshaw, Jukka Hakkola, Kari I. Kivirikko, Johanna Myllyharju, Raisa Serpi, and Peppi Koivunen. 2016. “Hypoxia-Inducible Factor Prolyl 4-Hydroxylase-2 Inhibition Protects Against Development of Atherosclerosis.” Arteriosclerosis, Thrombosis, and Vascular Biology. doi: 10.1161/ATVBAHA.115.307136.

Santolini, Marc, and Albert László Barabási. 2018. “Predicting Perturbation Patterns from the Topology of Biological Networks.” Proceedings of the National Academy of Sciences of the United States of America 115(27):E6375–83. doi: 10.1073/pnas.1720589115.

Schächinger, Volker, Martina B. Britten, and Andreas M. Zeiher. 2000. Prognostic Impact of Coronary Vasodilator Dysfunction on Adverse Long-Term Outcome of Coronary Heart Disease.

Schneider, Caroline A., Wayne S. Rasband, and Kevin W. Eliceiri. 2012. “NIH Image to ImageJ: 25 Years of Image Analysis.” Nature Methods.

Shi, Fanchao, Luanluan Sun, and Stephen Kaptoge. 2021. “Association of Beta-2-Microglobulin and Cardiovascular Events and Mortality: A Systematic Review and Meta-Analysis.” Atherosclerosis 320:70–78. doi: 10.1016/j.atherosclerosis.2021.01.018.

Souilhol, Celine, Jovana Serbanovic-Canic, Maria Fragiadaki, Timothy J. Chico, Victoria Ridger, Hannah Roddie, and Paul C. Evans. 2020. “Endothelial Responses to Shear Stress in Atherosclerosis: A Novel Role for Developmental Genes.” Nature Reviews Cardiology 17(1):52– 63.

Springstead, James R., B. Gabriel Gugiu, Sangderk Lee, Seung Cha, Andrew D. Watson, and Judith A. Berliner. 2012. “Evidence for the Importance of OxPAPC Interaction with Cysteines in Regulating Endothelial Cell Function.” Journal of Lipid Research 53(7):1304–15. doi: 10.1194/jlr.m025320.

Stancu, Camélia, and Anca Sima. 2001. “Statins: Mechanism of Action and Effects.” Journal of Cellular and Molecular Medicine. doi: 10.1111/j.1582-4934.2001.tb00172.x.

Steinberg, Daniel, and Joseph L. Witztum. 2010. “History of Discovery: Oxidized Low-Density Lipoprotein and Atherosclerosis.” Arteriosclerosis, Thrombosis, and Vascular Biology 30(12):2311–16.

Sterling, Teague, and John J. Irwin. 2015. “ZINC 15 - Ligand Discovery for Everyone.” Journal of Chemical Information and Modeling 55(11):2324–37. doi: 10.1021/acs.jcim.5b00559.

Sucosky, Philippe, Kartik Balachandran, Adnan Elhammali, Hanjoong Jo, and Ajit P. Yoganathan. 2009. “Altered Shear Stress Stimulates Upregulation of Endothelial VCAM-1 and ICAM-1 in a BMP-4- and TGF-?1-Dependent Pathway.” Arteriosclerosis, Thrombosis, and Vascular Biology. doi: 10.1161/ATVBAHA.108.176347.

Surdo, Prisca lo, Alberto Calderone, Gianni Cesareni, and Livia Perfetto. 2017. “SIGNOR: A Database of Causal Relationships between Biological Entities-a Short Guide to Searching and Browsing.” Current Protocols in Bioinformatics. doi: 10.1002/cpbi.28.

Tanaka, Tetsuhiro, and Kai Uwe Eckardt. 2018. “HIF Activation Against CVD in CKD: Novel Treatment Opportunities.” Seminars in Nephrology 38(3):267–76.

Theodoris, Christina v., Ping Zhou, Lei Liu, Yu Zhang, Tomohiro Nishino, Yu Huang, Aleksandra Kostina, Sanjeev S. Ranade, Casey A. Gifford, Vladimir Uspenskiy, Anna Malaschicheva, Sheng Ding, and Deepak Srivastava. 2020. “Network-Based Screen in IPSC-Derived Cells Reveals Therapeutic Candidate for Heart Valve Disease.” Science 0724(February):eabd0724. doi: 10.1126/science.abd0724.

Tong, Hanghang, Christos Faloutsos, and Jia Yu Pan. 2008. “Random Walk with Restart: Fast Solutions and Applications.” Knowledge and Information Systems 14(3):327–46. doi: 10.1007/s10115-007-0094-2.

Türei, Dénes, Tamás Korcsmáros, and Julio Saez-Rodriguez. 2016. “OmniPath: Guidelines and Gateway for Literature-Curated Signaling Pathway Resources.” Nature Methods.

Vogler, Melanie, Anke Zieseniss, Amke R. Hesse, Elif Levent, Malte Tiburcy, Eva Heinze, Nicolai Burzlaff, Gunnar Schley, Kai Uwe Eckardt, Carsten Willam, and Dörthe M. Katschinski. 2015. “Pre-and Post-Conditional Inhibition of Prolyl-4-Hydroxylase Domain Enzymes Protects the Heart from an Ischemic Insult.” Pflugers Archiv European Journal of Physiology. doi: 10.1007/s00424-014-1667-z.

de Vries, Margreet R., and Paul H. A. Quax. 2016. “Plaque Angiogenesis and Its Relation to Inflammation and Atherosclerotic Plaque Destabilization.” Current Opinion in Lipidology.

Willighagen, Egon L., Andra Waagmeester, Ola Spjuth, Peter Ansell, Antony J. Williams, Valery Tkachenko, Janna Hastings, Bin Chen, and David J. Wild. 2013. “The ChEMBL Database as Linked Open Data.” Journal of Cheminformatics. doi: 10.1186/1758-2946-5-23.

Wirka, Robert C., Dhananjay Wagh, David T. Paik, Milos Pjanic, Trieu Nguyen, Clint L. Miller, Ramen Kundu, Manabu Nagao, John Coller, Tiffany K. Koyano, Robyn Fong, Y. Joseph Woo, Boxiang Liu, Stephen B. Montgomery, Joseph C. Wu, Kuixi Zhu, Rui Chang, Melissa Alamprese, Michelle D. Tallquist, Juyong B. Kim, and Thomas Quertermous. 2019. “Atheroprotective Roles of Smooth Muscle Cell Phenotypic Modulation and the TCF21 Disease Gene as Revealed by Single-Cell Analysis.” Nature Medicine 25(8):1280–89. doi: 10.1038/s41591-019-0512-5.

Wu, David, Ru-Ting Huang, Robert B. Hamanaka, Matt Krause, Myung-Jin Oh, Cheng-Hsiang Kuo, Recep Nigdelioglu, Angelo Y. Meliton, Leah Witt, Guohao Dai, Nanduri R. Prabhakar, Yun Fang, Gö Khan, and M. Mutlu. 2017. “HIF-1a Is Required for Disturbed Flow-Induced Metabolic Reprogramming in Human and Porcine Vascular Endothelium.” doi: 10.7554/eLife.25217.001.

Xie, Liang, Xinchun Pi, Ashutosh Mishra, Guohua Fong, Junmin Peng, and Cam Patterson. 2012. “PHD3-Dependent Hydroxylation of HCLK2 Promotes the DNA Damage Response.” Journal of Clinical Investigation. doi: 10.1172/JCI62374.

Xu, Suowen, Iqra Ilyas, Peter J. Little, Hong Li, Danielle Kamato, Xueying Zheng, Sihui Luo, Zhuoming Li, Peiqing Liu, Jihong Han, Ian C. Harding, Eno E. Ebong, Scott J. Cameron, Alastair G. Stewart, and Jianping Weng. 2021. “Endothelial Dysfunction in Atherosclerotic Cardiovascular Diseases and Beyond: From Mechanism to Pharmacotherapies.” Pharmacological Reviews 73(3):924–67. doi: 10.1124/pharmrev.120.000096.

Yu, Guangchuang, Li Gen Wang, Yanyan Han, and Qing Yu He. 2012. “ClusterProfiler: An R Package for Comparing Biological Themes among Gene Clusters.” OMICS A Journal of Integrative Biology. doi: 10.1089/omi.2011.0118.

Yu, Guangchuang, Li Gen Wang, and Qing Yu He. 2015. “ChIP Seeker: An R/Bioconductor Package for ChIP Peak Annotation, Comparison and Visualization.” Bioinformatics 31(14):2382–83. doi: 10.1093/bioinformatics/btv145.

Zaffaroni, Gaia, Satoshi Okawa, Manuel Morales-Ruiz, and Antonio del Sol. 2019. “An Integrative Method to Predict Signalling Perturbations for Cellular Transitions.” Nucleic Acids Research 47(12):e72–e72. doi: 10.1093/nar/gkz232.

Zaffaroni, Gaia, Satoshi Okawa, Manuel Morales-Ruiz, and Antonio del Sol. 2019. “An Integrative Method to Predict Signalling Perturbations for Cellular Transitions.” Nucleic Acids Research 47(12):E72. doi: 10.1093/nar/gkz232.

Zhu, Lihua J., Claude Gazin, Nathan D. Lawson, Hervé Pagès, Simon M. Lin, David S. Lapointe, and Michael R. Green. 2010. ChIPpeakAnno: A Bioconductor Package to Annotate ChIP-Seq and ChIP-Chip Data.

Zhu, Tian, Shuyi Cao, Pin Chih Su, Ram Patel, Darshan Shah, Heta B. Chokshi, Richard Szukala, Michael E. Johnson, and Kirk E. Hevener. 2013. “Hit Identification and Optimization in Virtual Screening: Practical Recommendations Based on a Critical Literature Analysis.” Journal of Medicinal Chemistry.

